# Dopamine selectively regulates pediatric sclera/choroid interactions through the stimulation of exosome-associated retinoic acid

**DOI:** 10.1101/2025.07.13.664562

**Authors:** P Drugachenok, Y Ren, T Bai, L Rotard, A Dahlmann-Noor, M Bailly

**Author notes:** These authors have contributed equally (Y Ren, T Bai).

## Abstract

Postnatal eye growth is critical for healthy vision and yet, how this process is regulated at the cellular level is still unclear. The choroid is thought to play a crucial role in relaying signals from the retina to the sclera to modulate eye growth, but which cells are targeted and how they interact is not known. Using primary cultures of pediatric and adult human choroid and sclera stromal fibroblasts, we investigated the effect of choroid-conditioned medium (CCM) on scleral fibroblasts contractile activity. We show that CCM activates pediatric scleral fibroblast contraction, with age and antero-posterior location differences. Upon exposure to dopamine, a known negative regulator of eye growth, pediatric - but not adult - choroid cells lose their ability to stimulate scleral fibroblasts. Using RNA-Sequencing, we show that dopamine stimulates pathways linked to ribosomal activation, translation and exosome release exclusively in pediatric choroid cells. Removing exosomes from the CCM rescued the ability of dopamine-treated choroid cells to stimulate scleral fibroblasts. We further identify retinoic acid as the active exosome-associated compound preventing scleral fibroblast activation. Mechanistically, we show that CCM stimulates actomyosin-mediated protrusive activity in scleral fibroblasts. This is prevented when scleral cells are exposed to dopamine-CCM, and fully rescued upon exosome removal or ALDH1 inhibition. We thus propose that pediatric choroid stromal cells specifically relay dopamine-mediated retinal signals to the sclera during post-natal eye growth, operating through changes in secretome - including the production of exosome-associated retinoic acid - rather than gene expression changes.

**SIGNIFICANCE:** Eye size is critical to optimal vision but the exact cellular and molecular mechanisms regulating it are still unknown. Failure to properly regulate postnatal eye growth leads to elongated eyes and myopia. Myopia is expected to affect half of the world population by 2050, with many at risk of blinding complications. Eye growth and homeostasis are supported by the sclera and the choroid, but how the two tissues interact to regulate eye growth is unknown. We use here human cells to demonstrate direct, age-specific, interactions between the two cell types and provide a rationale for the known negative effect of dopamine on eye growth, with significant implications for future studies as well as for the understanding and treatment of myopia.

## INTRODUCTION

Eye size is a critical component of healthy vision. At birth, vision is imperfect, with a refractive defect called hyperopia: the eyes are too short for the light to come into focus on the retina, images are focused behind the retina, and vision is blurred. What happens over the course of the next 10 years or so is an exquisitely regulated growth process, balancing positive and negative influxes to achieve the state of emmetropia, i.e. where light is perfectly focused on the retina and vision is crisp. While it is clear that postnatal growth is predominantly regulated by light, the underlying cellular and molecular mechanisms have yet to be elucidated.

Eye growth and homeostasis are supported by the sclera and the choroid. From birth, the eye grows rapidly up until the age of 10, followed by a slow phase of minimal growth during adolescence^1^. This growth is supported by the expansion and maturation of the sclera, and upon reaching adulthood, the eye shape and size are irreversibly set by the final scleral shape and structure. The mammalian sclera is a stiff fibrous layer, composed mostly of collagen and proteoglycans, and within which reside cells, primarily fibroblasts. It is thought that cell proliferation and remodeling of the scleral extracellular matrix by the fibroblasts are required to allow eyeball growth and maturation, but the underlying mechanisms are unclear^2,3^. How scleral growth is regulated is still unknown.

The most recognized signal influencing eye growth is provided by light: light of high intensity, typically sunlight, promotes dopamine release in the retina. This in turn is thought to modulate eye growth by inhibiting sclera remodeling and axial elongation, through an as yet unidentified retina-to-sclera signaling cascade^3,4^. Although the relay secondary signals are unknown, evidence from animal models suggests that the choroid, the complex vascular-rich tissue lying between the retina and the sclera, may be central to this cascade, and to the overall maintenance of scleral homeostasis^3–6^. Choroid thickness changes during visual manipulations, thinning during accommodation and myopia^4,7^ and becoming thicker when recovering from light deprivation, and these changes are accompanied by a modified secretome^8,5^.

Of the many pathways, growth factors and molecules that have been implicated in eye growth, dopamine and retinoic acid (RA) both appear to be linked to a choroid effect on scleral growth. Dopamine is a critical modulator of eye growth, secreted in the retina upon light exposure. Dopamine can diffuse to the choroid, where it has been implicated in negative signaling to the sclera^4,9^. The role of retinoic acid appears more complex. Systemic or prolong dietary treatment with exogenous retinoic acid increases eye growth and promotes myopia in mice, guinea pigs and chicks^10,11^ ^12^. This is correlated with elevated retinoic acid levels in the retina (*reviewed in*^9^). However, in chicks, the levels of retinoic acid in the choroid appear to fluctuate in opposite direction: ocular elongation is linked a decrease in RA, while RA is increased in conditions that lead to decreased rates of ocular elongation^13^. The choroid was found to be the primary producer of RA, synthesizing and actively secreting considerably more RA than the retina, and all-trans retinoic acid (atRA) was shown to inhibit scleral proteoglycan synthesis, thus potentially directly regulating sclera growth and remodeling^13–15^. Furthermore, in chick eyes recovering from accelerated growth following form deprivation, both the mRNA levels of the main RA-producing enzyme retinaldehyde dehydrogenase 2 (RALDH2, also known as ALDH1A2) and the levels of atRA were elevated compared to paired controls^15^, suggesting that choroid RA might be specifically linked to signals leading to inhibition of eye growth/remodeling. Puzzlingly, oral delivery of RA in chicks induced eye elongation similarly to the other animal models, suggesting that exogenous untargeted RA delivery might act in a different manner compared to local physiological RA production^12^. Although the opposite eye growth response to RA in the chick compared with other animal models has been partly explained by the special conformation of the chick sclera (which includes a cartilage layer absent from mammal eyes^16^), the accompanying opposite changes in RA levels in the choroid remain unexplained, suggesting that the effects of choroidal RA on scleral growth might be species-specific^17^.

Furthermore, although RALDH2 positive cells have been found in the human choroid^5^, most of the data so far has been acquired from animal models where vision has been artificially manipulated, and there is little evidence that any of these mechanisms are directly at play during normal human postnatal eye growth.

We have used here human primary scleral and choroid cells derived from eyes of pediatric and adult donors to understand how the two cell types interact. We hypothesized that during postnatal eye growth, the choroid stroma integrates positive and negative signals to regulate sclera homeostasis though the release of soluble mediators, and that sclera/choroid interactions might evolve with postnatal age, dictating the course of postnatal eye growth and maturation. We used *in vitro* assays to demonstrate that pediatric choroid stromal cells specifically stimulate scleral fibroblasts through the release of soluble mediators, and that this stimulation is prevented when the choroid cells are treated with dopamine, the main negative regulator of eye growth. We further uncovered that the dopamine negative input is mediated through the release by pediatric choroid stromal cells of exosome-associated retinoic acid.

## RESULTS

### Dopamine modulates sclera/choroid stromal cells interactions

The primary tissue setting eye size and shape is the sclera^3^. The sclera is spatially compartmented, with noticeable antero-posterior differences in terms of matrix composition and stiffness^2,3^ and, significantly, the growing sclera of children is thinner and softer than adult sclera^18^, suggesting that the tissue changes with age in terms of composition and mechanical properties. The choroid cellular composition also is thought to evolve dynamically with age^10^. Specifically, the choroid stroma was shown to harbor multiple fibroblast-like subpopulations, including canonical fibroblasts, mural cells and non vascular smooth muscle cells (NVSMCs), which have been proposed to be involved in the regulation of the choroid function during eye growth and maturation (both in mouse and human)^19,20,21,22,23^.

We postulated that choroid stromal cells are central mediators in the retina-to-sclera signaling pathway driving post-natal eye growth, and that these cells might directly affect scleral fibroblast behavior and function through the release of soluble mediators.

Furthermore, we hypothesized that scleral fibroblasts – choroid stroma interactions change with age to regulate postnatal eye growth. To test these hypotheses, we cultured isogenic primary fibroblastic populations from the sclera and choroid of pediatric (3-7 year old) and adult (31-42) donors with no prior reported ocular issue. Scleral fibroblasts were obtained separately from both the anterior and posterior sections of the eyes, to account for the previously reported differences between anterior and posterior scleral tissue^2,3^. Mixed fibroblastic cell populations were obtained from whole choroids, following rapid growth from small explants. The primary choroid stromal cell cultures homogeneously expressed vimentin, with a subset expressing smooth muscle cell markers alpha smooth muscle actin (a-SMA), calponin, or both, suggesting the presence of a population of non-vascular smooth muscle cell (Supplementary Figure S1), as previously reported^19,22,23^. They were negative for pericyte markers NG2 and PDGFR, as well as endothelial cell markers CD31, CD144 and VWF (not shown), confirming the overall fibroblastic phenotype.

We first looked at the ability of choroid conditioned medium to stimulate scleral fibroblasts, and used cell-mediated collagen gel contraction as a proxy for general cytoskeletal and overall cell activation^24–26^. Paediatric scleral fibroblasts differ in their baseline ability to contract collagen under no stimulation (control serum free medium, SFM), with cells originating from the anterior sclera showing significantly more intrinsic force^24^ (Fig. 1A). Both anterior and posterior scleral fibroblasts showed increased contractile activity when exposed to paediatric choroid conditioned medium (CCM), and a milder response when exposed to adult CCM (Fig.1A). Posterior cells overall showed the greatest sensitivity to choroid-conditioned medium, particularly paediatric-CCM, with an approximate 2.5 fold increase in contraction potential when exposed to paediatric –CCM compared to SFM (1.6 fold for Adult-CCM), much greater than the effect observed in anterior cells (about 1.3 fold for both paediatric and adult CCM, Supplementary figure 2A). Overall this showed that human choroid stromal cells secrete compounds that can activate scleral fibroblasts, and that the interactions between the two cell types are more prominent with primary cells isolated from paediatric donors. Furthermore, we confirmed that scleral fibroblasts are heterogeneous with regards to antero-posterior spatial distribution, with cells isolated from the posterior sclera being less contractile but more responsive to stimulation compared to isogenic anterior scleral cells, consistent with a less mature growth stage^27^. Because of this and the fact that the posterior sclera is physiologically more likely to be regulated (as scleral changes associated with myopia and eye growth are more prominent at the back of the eye^27,28^), we focused on paediatric posterior scleral fibroblasts for the rest of this study.

**Figure 1.**
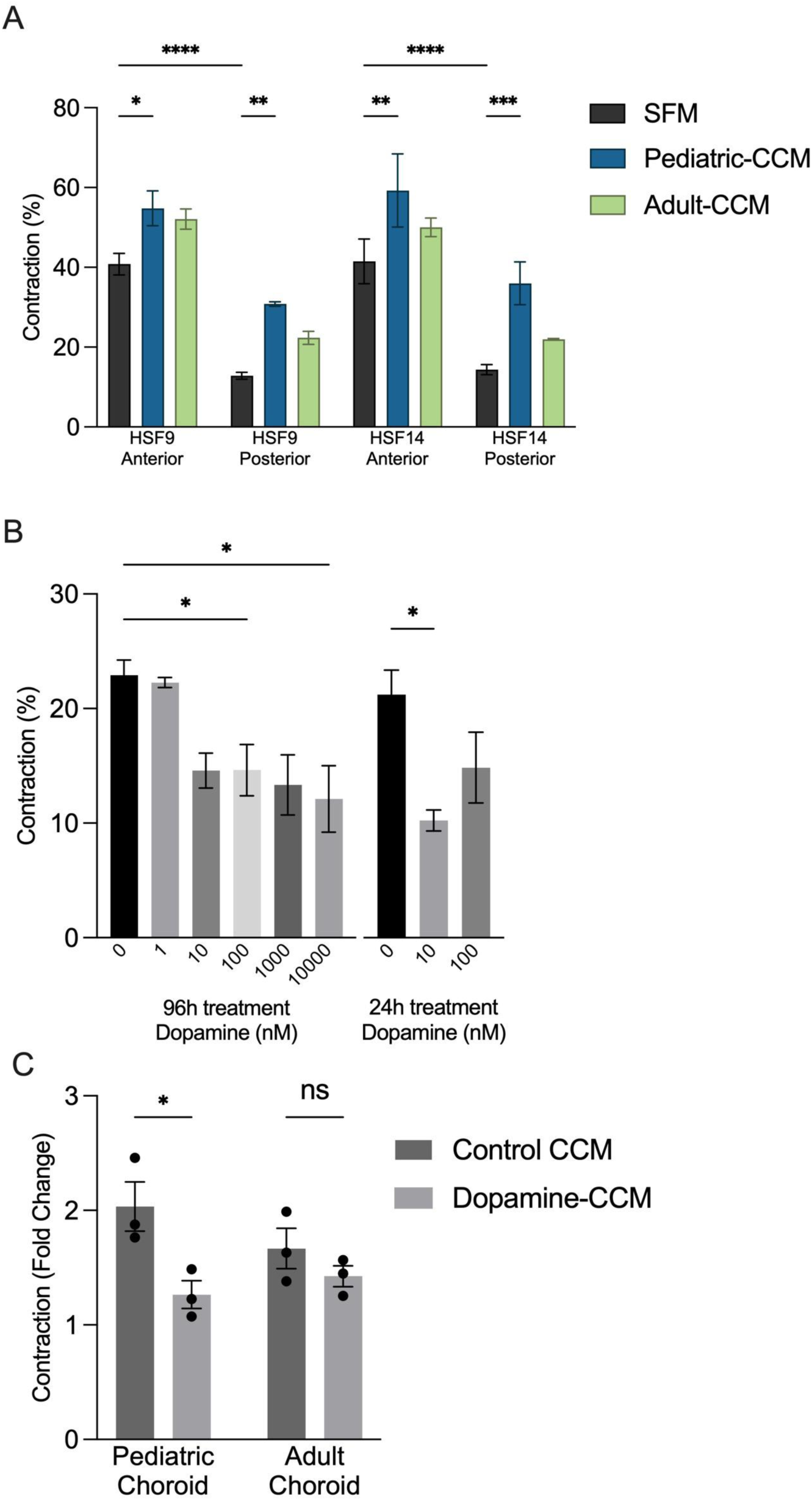
Dopamine treatment counteracts choroid cells’ ability to stimulate pediatric scleral fibroblast contractility. *(A)* Scleral fibroblasts from anterior and posterior sclera from two paediatric donors were embedded in collagen gels and contraction was measured after 1 day in the presence of the serum-free medium (SFM) or choroid-conditioned medium (CCM) prepared from paediatric (Choro-9R; Pediatric-CCM) or adult (Choro-10R; Adult-CCM) choroid cells. Shown is mean contraction (%) +/- SEM (SFM, n=6; each CCM, n=3). *(B)* Paediatric choroid cells (Choro9R) were treated with dopamine (1nM -10uM) for 96 or 24 hrs, after which the medium was then replaced with the serum-free medium for 24 hours to generate CCM and Dopamine-CCM. The CCM and Dopamine-CCM were used to stimulate paediatric scleral fibroblasts (HSF9-posterior) contraction. Shown is mean contraction (%) +/- SEM, n>=3. *(C)* CCM and Dopamine-CCM (10nM dopamine for 24 hours) were prepared from 3 paediatric (3, 3 and 7 year old) and 3 adult (31, 35, 42 years old) choroid cell lines, and used to stimulate paediatric scleral fibroblasts contraction (HSF9-posterior). Shown is mean contraction (%) +/- SEM (each point is the average of 3 independent experiment for each donor). * P<0.05, ** P<001, *** P<0.001, ****P<0.0001 (One- or Two-way ANOVA and Tukey’s multiple comparison tests).

Dopamine is a prominent regulator of postnatal eye growth. It is primarily secreted by the retina, including RPE cells^9,29^, and choroid cells have been proposed to be sensitive to dopamine^9,30,31^. We thus next assessed whether dopamine could act directly on choroid stromal cells to modulate their ability to stimulate scleral fibroblasts. We first treated paediatric choroid stromal cells (Choro-9, 3 year old donor) with a range of sub-physiological dopamine concentrations for 96 hours, then replaced the medium with dopamine-free serum-free medium to prepare CCM as before, and analysed the effect of the dopamine treatment on the contraction of isogenic paediatric posterior scleral fibroblasts (HSF9-Posterior).

Dopamine treatment had no visible effect on choroidal stromal cells morphology, and only minimal toxicity after 96 hours treatment, even with up to 1 uM dopamine (Supplementary Fig. S2 C, F). Treatment with dopamine concentrations as low as 10nM fully inhibited the ability of the paediatric CCM to stimulate scleral fibroblast contraction, bringing the contraction back down to the baseline levels seen in the presence of control serum free medium (Fig. 1B). This was not seen with adult choroid stromal cells (Choro-10, 35 year old), where even 10 uM dopamine could not fully prevent CCM-mediated paediatric scleral fibroblast contraction (Supplementary Fig. S2 B). Further work confirmed that only 24 hours incubation with 10 nM dopamine was sufficient to negate the choroid stromal cells’ ability to stimulate paediatric scleral fibroblast contraction (Fig. 1B), and these conditions were kept for all further experiments. The dopamine-CCM did not significantly affect scleral fibroblast metabolic activity/proliferation (as measured using CCK8 assay, Supplementary Fig. S2 D), total cell numbers (direct cell counts, Supplementary Fig. S2 E), or cell viability in the gels after 1 day (as measured using live/dead assay; Supplementary Fig. S2 G). Expanding the study to 3 paediatric (3, 3 and 7 year old) and 3 adult (31, 35, and 42 years old) choroid stromal cell cultures, we demonstrated that while paediatric choroid cells were only slightly better than adult ones at stimulating paediatric fibroblasts contracting ability, they displayed a unique intrinsic sensitivity to dopamine, which resulted in the abolishment of their stimulatory activity towards scleral fibroblasts (Fig. 1C). Adult choroid cells appeared more or less insensitive to dopamine in that respect.

### The effect of dopamine on choroid-conditioned medium is linked to translational changes and exosome production

To better understand the differences between paediatric and adult choroid stroma, and identify molecular candidates underlying the response to dopamine, we performed bulk RNA sequencing (RNA-seq) analysis on 3 paediatric and 3 adult choroid stromal cultures untreated (control) or treated with 10 nM dopamine for 24 hours. Normalization and differential analysis were carried out according to the DESeq2 model and package, comparing pediatric and adult, and control versus dopamine (full data set in Supplementary data file 1). The overall profiles showed little or no expression of endothelial marker CD144 (*CDH5*), CD31 (*PECAM*) and VWF (*VWF*), confirming the fibroblastic nature of the cells, with low levels of expression of pericyte markers NG2 (*CSPG4*) and PDGFR (*PDGFRA, PDGFRB*). Typical fibroblast markers were expressed at high level, including *VIM*, *COL1A1*, *THY1* and *DCN*, as were mural cell (smooth muscle cells and pericytes) markers ACTA2, CNN1-3, RGS4, RGS5, DES, and LMOD1^10,19,32,33^. Furthermore, both paediatric and adult stromal cultures expressed markers of mesenchymal stromal cell phenotype as previously described^34^ (CD34/*CD34* -, CD45/*PTPRC* -, HLA-DR/*HLA-DRB1*-, CD73/*NT5E*+, CD90/*THY1* +, and CD105/*ENG*+). Comparisons between paediatric and adult and with/without dopamine treatment were performed following fdr p-value adjustment, and a level of controlled false positive rate set to 0.05 (Fig. 2A, B). Consistent with the lack of effect of dopamine on adult CCM’s ability to stimulate scleral fibroblasts (Fig. 1C), we did not detect changes in gene expression following dopamine treatment of adult cells (Fig. 2A and Supplementary data file). Only 9 RNAs were significantly upregulated in dopamine-treated paediatric choroid stromal cells compared to control ones (3-7 fold change [log2]), none of them being actual protein genes: 3 pseudogenes (RNA5-8SP6, RNA5-8SP2, CTD-2328D6.1/DUXAP1), 5 RNA components of ribonucleoprotein complexes (RPPH1, RN7SL1, RN7SL2, RN7SK, RMRP) and a small nucleolar RNA (SNORD3A). These pointed out to global changes in ribosome biogenesis and translation, but did not give us direct insights on which component(s) change in the paediatric stromal cell secretome after dopamine treatment.

**Figure 2.**
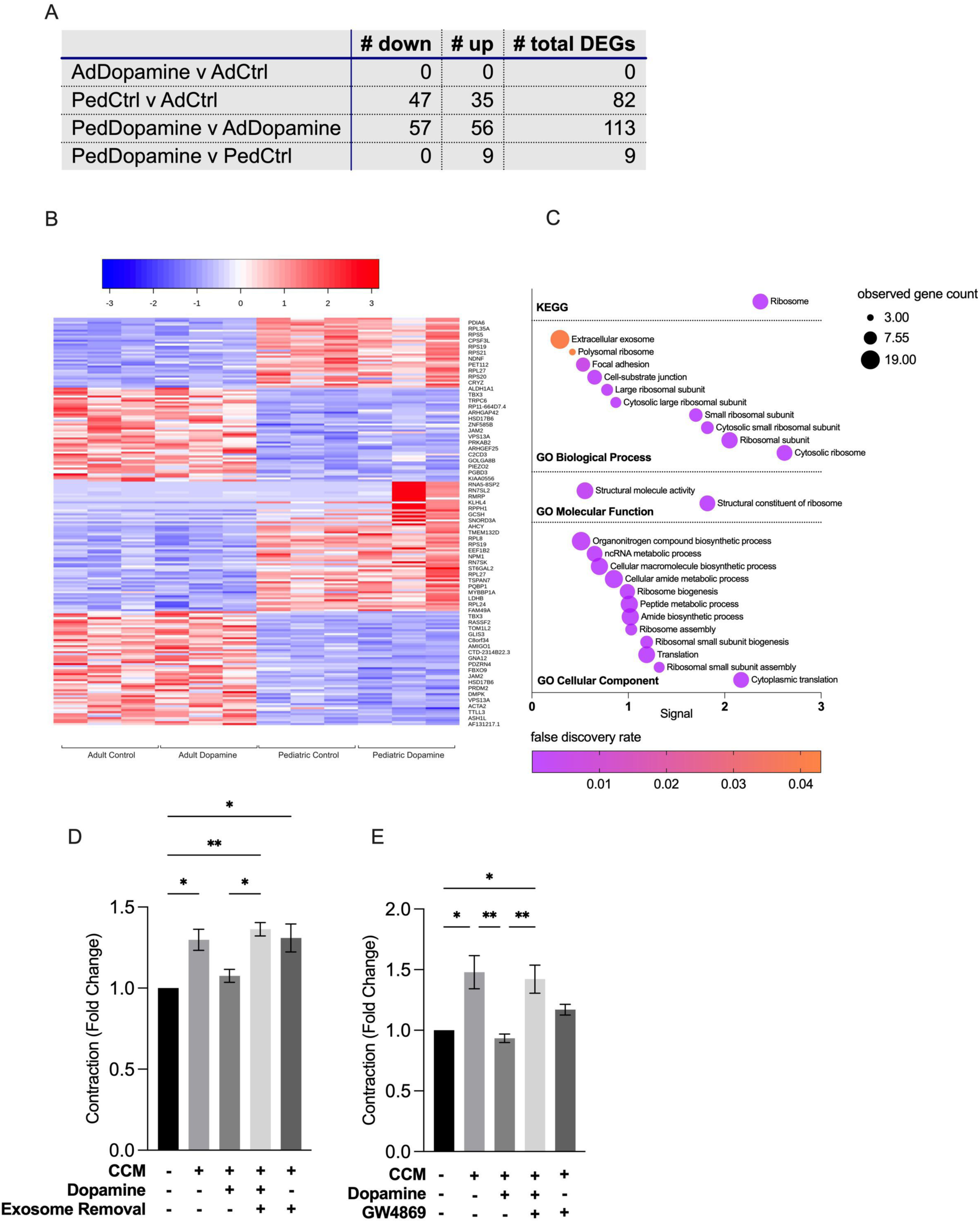
Ribosome/translational changes and exosome production mediate the effect of dopamine on choroid-conditioned medium. *(A)* Summary of differentially expressed genes (DGEs, adjusted fdr p-value < 0.05) comparing 3 paediatric and 3 adults choroid lines, with/without dopamine treatment (10 nM for 24 hours). *(B)* Heatmap visualization of DGEs showing the upregulation and downregulation of genes when comparing control and dopamine-treated paediatric choroid cells to control and dopamine-treated adult choroid cells, where each row represents a gene, and each column represents a sample. The samples are grouped into adult control, dopamine-treated adult, paediatric control and dopamine-treated paediatric cells. Within each row, the colour represents the fold change in normalised counts for each gene (blue, downregulation; red, upregulation in gene expression). *(C)* Pathway enrichment in DGEs comparing untreated (control) paediatric and adult choroid cells, as per KEGG and Gene Ontology pathway enrichment analyses. Where the signal (observed genes ratioed to background genes and normalised to false discovery rate) is represented on the x-axis, raw observed count is represented by the point size, and the false discovery rate is represented by the point colour. *(D)* CCM and Dopamine-CCM were prepared from paediatric choroid cells (Choro9L) and centrifuged x100000 g to pellet exosomes. Exosome-depleted CCMs and corresponding controls were used to stimulate scleral fibroblasts (HSF9-posterior) contraction. Shown as mean fold change in contraction (compared to serum free medium) +/- SEM (n=3). *(E)* Paediatric choroid cells (Choro9L) were treated with 10nM dopamine with/without 1µM GW4869 for 24hrs, after which the medium was replaced with serum-free medium for 24 hours to generate corresponding CCMs. These were used to stimulate scleral fibroblasts (HSF9-posterior) contraction. Shown as mean fold change in contraction (compared to serum free medium) +/- SEM (n=4). * P<0.05, ** P<0.01, as per One-way ANOVA and Tukey’s multiple comparisons tests.

To better understand the differences between paediatric and adult choroid stromal cells, we looked at the overall distribution of gene expression changes in the 4 different sets (Fig. 2B), and performed KEGG and Gene Ontology (GO) analysis on the full differential gene sets (FDR < 0.05) comparing paediatric control and adult control (Fig. 2C and Supplementary data file), and dopamine-treated paediatric cells versus dopamine-treated adult ones (Supplementary Fig. 3 and Supplementary data file). Both the KEGG and Gene Ontology analyses of genes differentially expressed between paediatric and adult choroid stromal cells overwhelmingly indicated an increase in RNA processing, ribosomal biogenesis and translation, suggesting that paediatric cells are more translationally active than adult ones (Fig. 2C; Supplementary data file). The same pathways and biological processes were dominant when comparing dopamine-treated paediatric to dopamine-treated adult (Supplementary Fig. 3, Supplementary data file). However, looking at the GO enriched cellular components, “extracellular exosomes” was the second most important after cytosol in terms of gene counts (Fig. 2C, Supplementary data file), suggesting that this increased ribosomal activity may lead to changes in exosome production. Overall these suggested that the dopamine-specific difference between paediatric and adult choroid stromal cells may not lie within direct changes in gene expression/mRNA levels but rather results from changes in translation and protein synthesis, as suggested by previous studies^35,36^, and possibly leading to increased release of exosome-associated contents.

To determine whether the inhibitory effect of dopamine-CCM on scleral fibroblast activation might be mediated by exosomes, we first use a standard ultracentrifugation protocol to deplete exosomes from dopamine-CCM prior to adding it to the scleral fibroblast contraction assay. Depleting exosomes from dopamine-CCM fully rescued the CCM’s ability to stimulate scleral fibroblast contraction (Fig. 2D). Similarly, co-treating choroid stromal cells with GW4869 to prevent exosome biogenesis/release during the dopamine incubation prevented the negative effect of dopamine on the CCM’s ability to stimulate paediatric scleral fibroblast contraction (Fig. 2E). These results suggested that dopamine triggers translational changes in paediatric choroid cells, which lead to release of exosomes, whose content is able to prevent scleral fibroblast activation.

### The dopamine-induced negative effect on scleral fibroblast activation is mediated by exosome-associated RA

RA is a longstanding candidate in the regulation of postnatal eye growth and myopia. The sclera is known to have retinoic acid receptors^37^, and RA is one of the few signals in the retina-to-sclera cascade that has been shown to have a direct effect on scleral cells^9,13^. RA is secreted primarily by the retina and the RPE, but there is also direct evidence that the choroid produces significant levels of RA^13^ ^38^. In particular, recent work using chick and human tissue has shown that a specific population of stromal choroid cells can produce RA^5^. Furthermore, some evidence suggests that functional RA can be released in association with exosomes^39^. We thus hypothesized that RA might be the exosome-associated inhibitory compound released by dopamine-treated choroid stromal cells that blocks scleral fibroblasts’ activation. We first measured RA content in the CCM and in exosomes from dopamine-CCM. Under normal culture conditions, the CCM produced by choroid paediatric stromal cells contains about 70 pM RA, with approximately 2/3^rd^ of it associated with the exosome fraction (Fig. 3A). Following treatment with 10nM dopamine for 24 hours, there was a 1.5 fold increase in the amount of RA measured (115 pM), again with the majority of it in the exosome fraction. We then assessed whether scleral contraction could be inhibited by RA. In the presence of 10% serum, paediatric scleral fibroblasts contracted collagen gels efficiently. The contraction was fully inhibited in the presence of 1uM RA, but rescued by addition of pan-retinoic acid receptor (RAR) antagonist AGN193109 (1uM), suggesting that RA acts on the scleral fibroblasts through activation of the RAR (Fig. 3B). Similarly, RA fully inhibited CCM-induced scleral fibroblasts’ contraction, and blocking RAR on the scleral fibroblasts fully restored the ability of dopamine-CCM to stimulate contraction, suggesting that the negative effect of the dopamine-CCM on scleral contraction is mediated by RA (Fig. 3C). To confirm this, we blocked RA synthesis in choroid stromal cells during their treatment with dopamine using ALDH1 inhibitor WIN18446, again fully restoring the CCM’s ability to stimulate scleral fibroblasts’ contraction (Fig. 3D). This overall suggested that paediatric choroid stromal cells by default produce a secretome that can activate scleral fibroblasts. Upon dopamine exposure for 24 hours, choroid stromal cells release exosome-associated RA that directly targets RAR in scleral cells and prevents activation and ensuing cell-mediated gel contraction.

**Figure 3:**
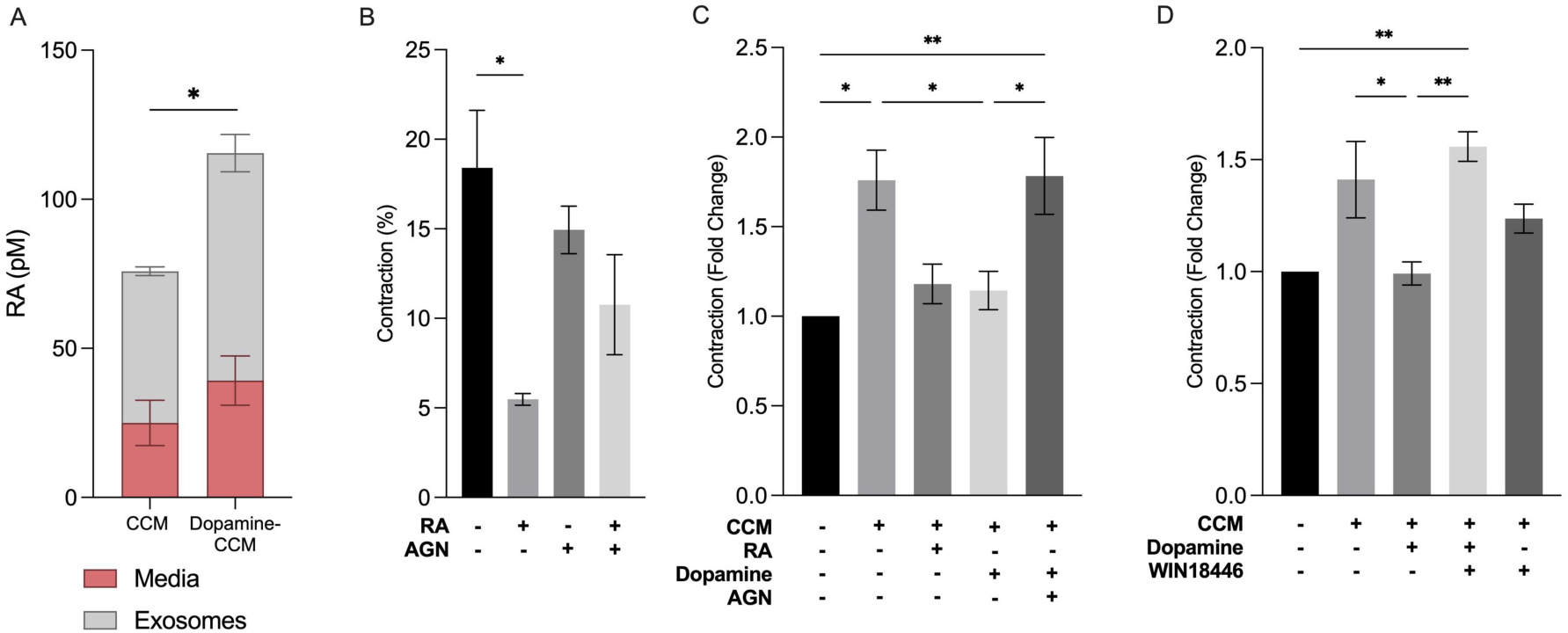
The dopamine-induced negative effect on scleral fibroblast activation is mediated by exosome-associated retinoic acid. *(A)* Retinoic acid concentration in the separated media and exosome fractions of CCM and Dopamine-CCM. Shown is RA concentration (pM) +/- SEM, n=3. Exosomal concentration was adjusted for full CCM volume. *(B)* Paediatric scleral fibroblast (HSF9-posterior) collagen gel contraction in serum containing medium (10% serum) with/without RA (1µM), RAR inhibitor AGN193109 (1µM) or both.Shown is mean contraction (%) +/- SEM, n=3. *(C)* CCM and dopamine-CCM were prepared from paediatric choroid cells and used to stimulate paediatric scleral fibroblast (HSF9-posterior) collagen gel contraction in the presence of RA (1µM), RAR inhibitor AGN193109 (1µM) or both. Shown is mean contraction (%) +/- SEM, n=3. *(D)* Paediatric choroid cells (Choro9L) were treated with 10nM dopamine with/without 8µM WIN18446 for 24hrs, after which the medium was replaced with the serum-free medium for 24 hours to generate CCM and Dopamine-CCM. These were used to stimulate scleral fibroblasts (HSF9-Posterior) contraction. Shown as mean fold change in contraction (compared to serum free medium) +/- SEM (n=4). * P<0.05, ** P<0.01, (One- or Two-way way ANOVA and Tukey’s multiple comparisons tests).

### Exosome-associated RA decreases scleral contraction through a reduction in acto-myosin contraction

We previously showed that fibroblast contraction activity in the free-floating collagen gels used in our assay is linked to intrinsic cellular force and cell protrusive activity, rather than proliferation^24,25,40^. Indeed, although RA significantly inhibited pediatric scleral fibroblasts proliferation in standard 2D cultures (Supplementary Fig. S4A), we did not observe significant proliferation in 3D collagen gels over the course of 7 days, nor was there an effect of RA, even at high concentration, despite the contraction being significantly affected (Supplementary Fig. S4 B, C). Using phalloidin staining to highlight overall cell shape, we thus analyzed scleral fibroblast morphological parameters following 1 day in culture with CCM, dopamine-CCM and dopamine-CCM with concurrent exosome biogenesis prevention (GW4869) or RA production inhibition (WIN18446). In the absence of stimulation (serum free medium, SFM) scleral fibroblasts in 3D gels remained fairly round, with few protrusions after 1 day of culture (Fig. 4A). Exposure to CCM triggered a dramatic change in cell shape, with the cells spreading extensively in the matrix and extending numerous protrusions. This was reflected in a significant increase in Aspect/Ratio and maximum length (as measured by Feret diameter, Fig. 4 A-C), and concomitant decrease in circularity and solidity (Fig. 4 D,E). Scleral fibroblasts exposed to dopamine-CCM remained rounder and with few protrusions, with morphological parameters identical to those of fibroblasts in serum free medium (Fig. 4, A-E). Preventing exosome biogenesis or RA production during dopamine treatment when preparing the CCM restored the scleral fibroblast spreading and all morphological parameters to levels similar to those exposed to untreated CCM, suggesting that exosome-associated RA produced by dopamine-treated choroid stromal cells directly affects the protrusive activity of scleral fibroblasts.

**Figure 4.**
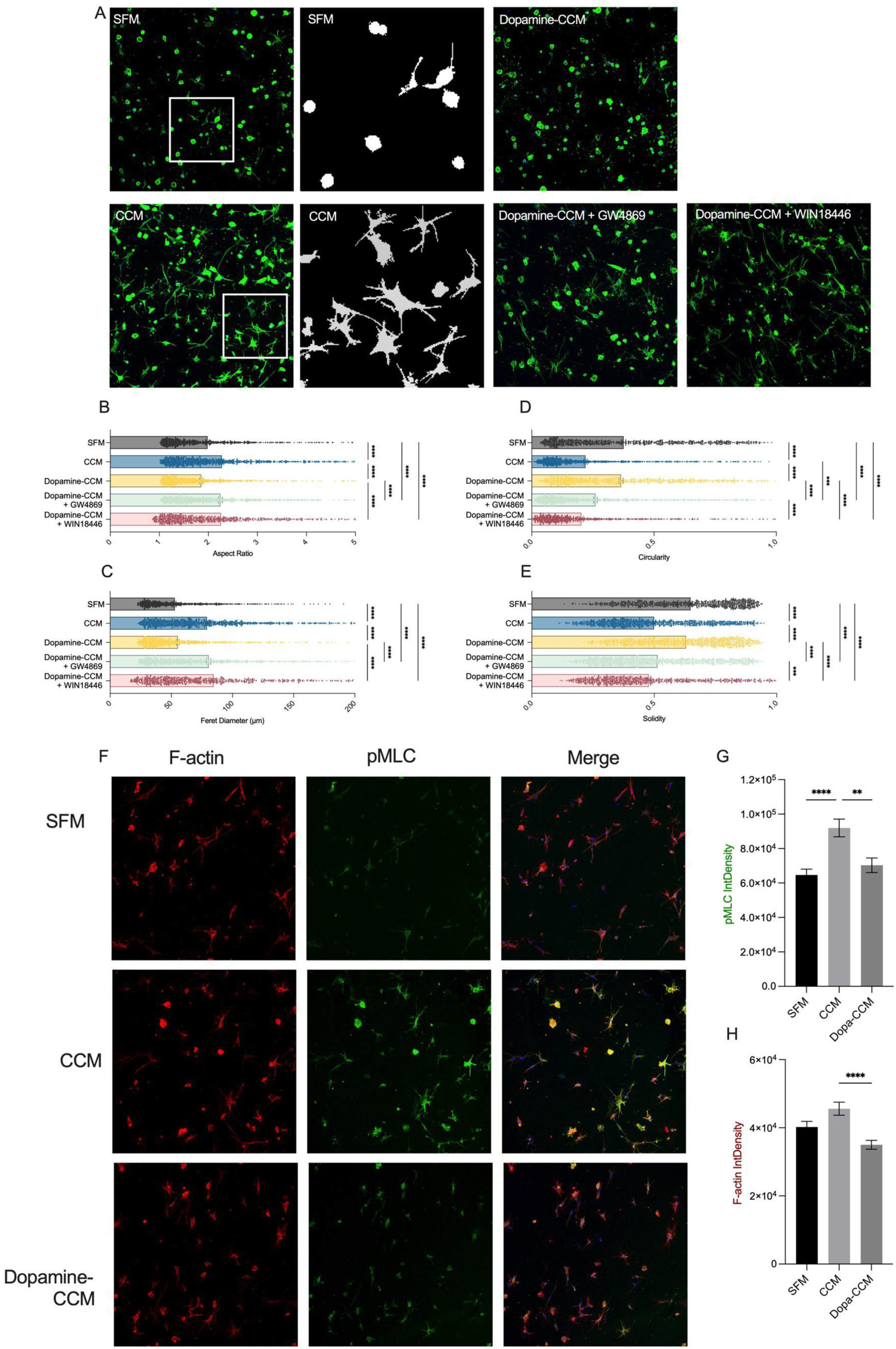
Exosome-associated RA decreases scleral contraction through a reduction in acto-myosin contraction. *(A)* Representative images of paediatric scleral fibroblasts (HSF9-Posterior) in collagen gels after one day in SFM, CCM, dopamine-CCM or dopamine-CCM obtained after treatment with GW4869 (1µM) or WIN18446 (8µM). Shown are z-projections of confocal stacks of images of cells stained with FITC-phalloidin. Inset shows typical masks used for measurements. Scale bar, 100 um. *(B-E)* Corresponding cell shape analysis comparing the aspect ratio *(B)*, Feret diameter *(C)*, Circularity *(D)* and solidity *(E)* of scleral fibroblasts. Each datapoint representing a cell (>200), from 3 independent repeats. *(F)* Representative images of paediatric scleral fibroblasts (HSF9-Posterior) in collagen gels after one day in SFM, CCM, dopamine-CCM and stained for phospho-myosin (green) and F-actin (red)*. (G, H)* Fluorescence quantitation for phospho-myosin *(G)* and F-actin *(H)*. Represented as average integrated density/cell +/- SEM (n=3). ** P<0.01, ****P<0.0001 (One-way ANOVA and Tukey’s multiple comparisons tests).

Protrusive activity is linked to dynamic changes in the cells’ cytoskeleton and acto-myosin contraction is most often a critical component of protrusive activity and contractile properties of fibroblasts in 3D matrices. Previous work suggested that RA derivatives could affect actomyosin contraction^41^ and myosin light chain (MLCII) phosphorylation^42^. To investigate possible changes in acto-myosin activity in gels, we evaluated the level of phosphorylation of myosin light chain (pMLC) using immunofluorescence. There was a significant increase in phospho-myosin in scleral fibroblasts stimulated with CCM compared to serum free medium medium (Fig. 4 F, G). This was completely abolished when choroid stromal cells were treated with dopamine prior to preparing the conditioned medium (dopamine-CCM). This was accompanied by a concomitant decrease in filamentous actin in the cells (Fig. 4 F, H), suggesting that the dopamine-CCM directly perturbed actin dynamics and acto-myosin networkss in scleral cells.

## DISCUSSION

Postnatal eye growth is a critical phase in healthy vision; it ensures the eye grows to a precise mature size where its structural parameters allow for incoming images to be perfectly focused on the retina, so called 20/20 vision or emmetropia^27^. Unlike other organs, the eye’s final size and maturation are not regulated centrally but depend on local signals, primarily through visual input: light signals coming to the retina (e.g. light intensity, blur level) are thought to be picked up by specific retinal populations, which in turn convert these vision signals into secreted biochemical messengers that trickle down from the retina to the sclera, in the so-called retina-to-sclera signaling cascade^9,27,6^. Although some of the major signals have been identified, including dopamine, RA and nitric oxide^6,9^, the actual cellular and molecular interactions linking inputs in the retina to an effect in the sclera still remain elusive^43^. In particular, while it is thought that the choroid plays a critical role in relaying the retinal signals to the sclera to modulate eye growth, we do not yet understand which cells are involved and what the exact signals are^7^. Furthermore, the vast majority of the research on postnatal eye growth has been done through visual manipulations in animal models, where eye growth is commonly modulated through form deprivation or the wearing of lenses^43^.

There is mounting evidence that the eye growth response to such drastic visual modifications might be very different from the day to day and circadian adjustments that occur during normal postnatal eye growth^9,27^, and that these animal models may not accurately reflect what’s happening in human eyes^44^. Finally, very little is know about the cellular composition of the human sclera and choroid with regards to postnatal changes, with only a handful of studies reporting analyses on paediatric tissue. To better understand postnatal eye growth and directly interrogate putative interactions between the choroid and the sclera, we have generated isogenic primary cultures of choroid and scleral stroma from donor eyes, comparing paediatric (growing) and adult (mature, stable) eyes and how the interactions between the two tissues are modulated with age.

We have identified here for the first time key interactions between stromal cells from the sclera and those of the choroid using our human model system. We showed that choroid cells secrete compounds that send a positive activation signal to scleral fibroblasts. These interactions are regulated by age, being more prominent in paediatric tissue, and by the cell location within the eye, with cells from the posterior sclera section being more receptive to choroid stimulation. We found that choroid stromal cell populations also change with age, with significant differences in gene expression between paediatric and adult cells, and that paediatric stromal cell populations are exclusively responsive to dopamine. Dopamine exposure appears to stimulate translational activity in paediatric choroid cells and results in the production of exosome-associated RA. In turn, these exosomes are able to fully counteract the baseline stimulatory activity of choroid cells. Our work thus suggests a previously uncovered age-specific mechanism by which dopamine regulates postnatal eye growth through stimulation of exosome release.

### Primary fibroblastic sclera and choroid cell populations retain intrinsic age and tissue specific characteristics

Fibroblast and stromal cell populations are known to be heterogeneous and dynamic with age and tissue location, with distinct cellular signatures underlying tissue homeostasis during growth and aging^45^. In the eye, we have shown previously that fibroblasts from different ocular locations present different functional properties, for example various level of intrinsic force that match the stiffness of their tissue of origin^24,25^. We also demonstrated that human primary ocular fibroblast cultures retain specific tissue state properties, particularly when cultured within more physiological 3D environments, and that they can be used to develop *in vitro* disease and tissue functional models^26,46–48^. We thus anticipated that sclera and choroid stromal cells might behave in a similar way. Indeed, we show here that paediatric choroid stromal cell populations are clearly different from adult ones, both in their native mRNA expression signatures as well as their functional response to dopamine stimulation. The cells derived from paediatric donors displayed a more transcriptionally active profile, with increased ribosome biogenesis and accompanying translation. This activity was further increased upon dopamine stimulation, while stromal cells from adult eyes displayed little or no response to dopamine.

Despite ample evidence suggesting a critical role for the sclera in supporting ocular tissue homeostasis, how the sclera is regulated during postnatal eye growth is unknown. The eye is thought to mature from front to back, with some evidence that the posterior sclera is different from that in the anterior section, resembling a “younger” tissue^27^. The sclera was shown to be spatially compartmented, with noticeable antero-posterior differences in terms of matrix composition, gene expression and overall stiffness^18,49,50,51^, as well as regional differences with regards to eye orientation (superior/lower and nasal/temporal^18,50,52^).

Significantly, the growing sclera of children is thinner than adult sclera, as well as noticeably softer, suggesting that the tissue also changes with age in terms of cellular composition and mechanical properties^18,51^. Consistent with this, we show here that anterior pediatric scleral fibroblasts contract collagen gels with much greater efficiency compared to isogenic posterior ones, reflecting a higher intrinsic force^24^ and predictive of a higher tissue stiffness *in vivo*^24,25^. This is consistent with a previous report suggesting that human anterior sclera is stiffer than the posterior section^51^. Furthermore, we found that posterior scleral cells were more sensitive to stimulation by choroid conditioned-medium than their anterior counterparts, reminiscent of a previous study that showed that the pediatric posterior sclera is more sensitive to activation of the Wnt/β-catenin signaling pathway, a highly conserved pathway that regulates key cellular functions including proliferation^49^. This suggests that the posterior pediatric sclera may be more receptive to signals from the choroid to regulate its expansion, and primary fibroblasts cultures retain some of this specific behaviour *in vitro*.

While the cellular profile of the human choroid has been investigated by many labs recently^8,19–21,32,33^, the majority of the studies used adult tissue, mostly in the context of AMD, and seldom focus on the stromal fibroblastic populations. Monavarfeshani et al. performed single cell sequencing of RPE/choroid on 8 adult donors (most of them over 60 years old) and identified two principal fibroblastic clusters (expressing *COL1A1* and *DCN*) in the choroid, with specific markers *TRDN* (C10 cluster) or *KCNQ3* and *LGR5* (C28 cluster)^32^.

Although both our paediatric and adult choroid stromal cell populations express *COL1A1*, and to a lesser extent *DCN*, *KCNQ3* and *LGR5* were expressed at very low level, and *TRDN* was not detected (Supplementary data file). This may reflect a lower frequency of these populations in primary cultures, or a differential expression of those markers with age, as the adult donors in our study are still significantly younger than those from Monavarfeshani et al. Thus, why some of the markers identified in fibroblastic clusters in adult human tissue were present in our stromal cell population, it appears that fibroblastic populations identified in adult choroid tissue may be significantly different from those in paediatric tissue. Consistent with our identification of smooth muscle cell markers in our primary stromal choroid populations, we found that, similar to cells in adult choroid tissue, they express mural cells/pericyte markers (cluster C26^32^), including *TAGLN* and *ACTA2*, as well as *TRPC4*, *RGS5*, and *LAMA2*. As for *COL1A1* and *DCN*, we found that most of these markers are also expressed at lower level in pediatric stromal cells compared to adult ones (Supplementary data file), suggesting that pediatric stromal choroid populations might be overall less differentiated. Collin et al. compared foetal and adult RPE/choroid cell populations^33^; although foetal cell populations may intuitively seem closer to paediatric cells than adult, they have not been exposed to the critical vision-guided phase of post-natal eye growth expected in paediatric cells. Nevertheless, our primary choroid stroma populations appear to express most of the markers expressed in clusters identified by Collin et al., with again most of them expressed at higher level in adult cells: neural crest cells (*PITX2 and FOXC1),* fibroblasts *(PENK, COL1A1, FBLN1*), pericytes *(THY1, ITGA1, POSTN*), smooth muscle cells *(ACTA2, TAGLN),* periocular mesenchyme *(MGP, DCN)* and nerve-associated fibroblast (*SFRP4, PI16*). Interestingly, 35/82 DEGs in our differential analysis comparing paediatric and adult stromal cells are present in the DEGs identified by Collin et al. when comparing foetal and adult cells in smooth muscle cell clusters (Supplementary data file 2), suggesting similarities between foetal and paediatric choroid populations, as well a pointing to a possible significant role of non-vascular smooth muscle cell populations. String analysis and functional enrichment of the 35 DGEs common between paediatric versus adult (our study) and foetal versus adult (Collin et al.^33^) again reveals a significant link to translation and ribosome biogenesis (Supplementary data file 2). This suggests that our primary choroid stroma cultures retain critical genomic expression signatures pertaining to the age and maturation of the tissue, and is consistent with previous work showing that ribosomal protein levels evolve with development stage and tissue specification^53^. An attempt at correlating gene expression changes with age was made by Huang et al, who performed single cell sequencing of RPE/choroid tissue from donors of different ages, including one foetal and one pediatric sample^10^. They found that the cellular composition of the choroid changed with age, with an increase in retrieved immune cells and a decrease in fibroblasts as the donor age increased. Interestingly, among the pathways and genes enriched with age in correlation analysis, they found “vesicle-mediated transport and its regulation” negatively correlated with age in the fibroblast populations identified (Supplementary table 5 in Huang et al.^10^), suggesting an increased involvement of these pathways in younger choroid tissues, consistent with our own study. Furthermore, “ribonucleoprotein complex biogenesis” and “ribonucleotide metabolic processes” were also negatively correlated with age in the pericyte cluster, again suggesting increased transcriptional processing in the younger cells (Supplementary table 5 in Huang et al.^10^), as we found in our study (with genes *RPL21*, *RPL27* and *RPL35* upregulated in pediatric versus adult in our study and negatively correlated with age in Huang et al.^10^).

Overall this suggests that human choroid stroma composition evolves with age, and appears to be significantly more transcriptionally active during the pre-and postnatal period, possibly reflecting a lower level of differentiation and maturation. Such an immature state is likely to make the pediatric choroid stroma more amenable to modulation through signals conveyed by visual clues to drive emmetropia.

### Pediatric, but not adult, choroid stromal cells are sensitive to dopamine

Dopamine has long been implicated in eye growth, presumably relaying light-induced retinal inputs to modulate eye growth^9,30,54^. However, the target cell(s) and the actual cellular effects are still unknown. Dopamine is primarily secreted in the retina, most likely by amacrine cells, but is also found in the choroid^31,54^. We show here for the first time that 10nM dopamine, a concentration consistent with the range of what can be estimated from animal models^55,56^, is sufficient to trigger a response from paediatric choroid stromal cells. Strikingly, we found that adult choroid cells were not sensitive to dopamine, confirmed by an absence of significant gene expression changes following an overnight stimulation with dopamine. This suggests for the first time that the postnatal period is a privileged period during eye growth, where possibly less differentiated choroid stromal cells are sensitive to vision-derived signals that are not picked up by the mature cells present in the adult eye. It is likely that the paediatric choroid would display greater sensitivity to positive signals driving eye growth, and in turn would more efficiently stimulate the sclera. Indeed, although both adult and paediatric choroid stromal cells are able to stimulate scleral contraction in our *in vitro* set up, there is a hint that the paediatric choroid stromal cells are indeed more efficient at stimulating paediatric scleral fibroblasts than are adult ones (at least when looking at posterior sclera, Supplementary Fig. S2 A). Further work will be needed to confirm this.

What exactly makes the choroid pediatric cells more responsive to dopamine is unclear. Aside from the major involvement of ribosomes/translational activity, the DGEs between pediatric and adult choroid cells do not point to obvious pathways that could link to a different response to dopamine (Supplementary data file). There is a hint that paediatric stromal cells have less of a smooth muscle phenotype compared to adult ones (actomyosin contraction [*CNN1*; *CALD1*; *ARHGAP42*; *PIEZO2*] and cAMP regulation [*IGFBP5*; *C8orf34*], components all down in paediatric versus adult control DGEs, Supplementary data file 1), and a more dynamic cytoskeleton (increase in a Rac GEF *PREX1*), with some evidence suggesting a more proliferative phenotype (negative regulation of cell division [*SUSD2*] is down in paediatric). This could suggest the presence of an immature proliferative non-vascular smooth muscle cell population in the paediatric choroid stroma^23^. Whether this is one population or multiple ones, and which one is responding to dopamine remains to be confirmed. In addition, a number of reports in the literature link dopamine stimulation and increased ribosomal/transcriptional activity in neurons^57,58,35,36^. We cannot exclude the possibility that our paediatric choroid stromal cells contain a population of dopamine-sensitive neurons that are responding acutely to the stimulation (indeed neuronal marker *SCN9A* and Schwann cell markers *PRNP* and *MBP*^19^ were found to be expressed at significant levels in both paediatric and adult choroid populations, Supplementary data file 1).

### Dopamine stimulates the release of exosome-associated RA

RA has long been thought to be involved in the regulation of eye growth^5,9,13^ and yet, similar to dopamine, its cellular targets and the resulting molecular consequences are unclear. RA, and RA-producing enzymes ALDH/RALDH are present in the choroid^5^, and RA levels in the choroid (and in the retina) are fluctuating in concert with visual manipulations and eye growth^9^. In the chick, at-RA content is reduced in the choroid during induction of myopia and increased during induction of hyperopia, with RALDH2/ALDH1A2 mRNA levels following the same pattern, reduced when eyes were exposed to negative lenses (increased eye growth) and increased following exposure to positive lenses (reduced eye growth)^15,59,60^. Although these early works suggested that RA in the choroid functions as a negative regulator of eye growth, exogenous treatment with RA was shown to cause excessive eye growth and myopia in chicken, mouse and guinea pigs^9^. Mc Fadden et al. argued that “RA may act at the level of a coordinated non-visual regulatory system which controls the growth of the various ocular components”^12^, while others suggested that the contribution of RA to normal growth might be different to that of myopia^27^ or visually manipulated growth^60^ and/or dependent on the age of the animal studied^11^ (effectively differentiating between young, growing eyes and eyes at the end of emmetropisation^12,17^). One major drawback of these studies is that they all look at a global effect of RA on tissue, without means to understand where and how RA acts. RA may target cells in the retina, in the choroid, and/or the sclera. These tissues contain multiple target populations that might react in different ways, or be sensitive to different levels of RA. It is entirely possible that RA’s function in one tissue (e.g. the retina) counteracts its function in another tissue (e.g sclera). Early work using the chick model found that the choroid produces large amounts of RA, which is actively secreted and can readily reach the sclera^13^. Similarly, in late adolescent and adult human eye, ALDH1 catalytic activity was detected in the choroid, at significantly higher levels than in the RPE or the retina^61^. RALDH2/ALDH1A2 and RALDH1/ALDH1A1 proteins were found associated with stromal cells in the human choroid^5,61^, consistent with the significant mRNA levels of the corresponding genes in our adult choroid stromal cell population (Supplementary data file 1). Of note, both genes are expressed at much lower levels in our paediatric choroid stromal cells set, suggesting that either RA synthesizing cells are in much lower numbers in the paediatric choroid stroma, or RA synthesis by paediatric choroid cells happens through a different pathway. The level of RA that we measured (around 100 pM) is significantly lower than what’s been measured or estimated in animal models, where choroidal RA was measured in the nanomolar range^13,15,62^. However, our conditioned medium is highly diluted compared to tissue, with a final volume of 8 ml for <10^6^ cells. We estimate that it is about 500 times less concentrated than tissue, which would give a concentration of 50nM for a tissue equivalent, well within the range of *in vivo* values that were shown to affect scleral proteoglycan synthesis^13,15^.

Chick choroids placed in tissue culture were shown to secrete RA with rapid kinetics, and after 30 min there was already more RA in the medium than in the cells suggesting very active secretion in a rather stable form, with most of the secreted RA eventually finding its way to the sclera. The authors found that choroid RA was secreted in complex with a 28-32 kD protein^13^, which is consistent with a more stable form of RA, including a possible association with exosomes (yet to be identified at the time). Interestingly, an association between RA and exosomes has been previously reported in oligodendrocytes precursors: the cells were shown to synthesize RA and release it in association with exosomes, in a process that regulates myelination in the central nervous system^39,63^. In recent years, numerous studies have identified exosomes in the eye, with potential links to pathology^64^.

While most reports look at RPE-derived exosomes^65^, exosome-like vesicles have been observed in donor eyes from patient with choroideremia, around Müller cells and throughout the retina and subretinal space^66^. In both human and chicken adult choroid, melanopsin and VIP immunostaining revealed a pattern of granules in the stroma reminiscent of extracellular vesicles^67^, and RALDH2 was similarly found in a granular pattern in human choroid stroma cells^5^. It is thus conceivable that exosomes may play a hitherto undiscovered yet significant role in the choroid. Releasing mediators of eye growth such as RA through exosomes may ensure that transient signals such as those provided by visual inputs perdure longer than the equivalent free biological molecule (growth factor, RA…) by regulating the release and stability of the exosome-associated compound as well as ensuring targeted delivery. On the other hand, the post-transcriptional rather than transcriptional regulation may ensure a more dynamic modulation of the response, which could account for how the postanal eye can integrate multiple, often contradictory, visual signals to fine-tune growth^27^.

### Dopamine-induced exosomes affect pediatric fibroblast dynamics in 3D environments

The effect of RA on sclera has largely focused on the production of GAGs, as a reduction in GAG production has long been associated with sclera remodeling and thinning during myopia and following visual manipulations leading to increased eye growth^9,13,38^.

However, the kinetics of changes in GAG production are often in the order of days, thus more likely linked to changes in transcriptional regulation, and the underlying signaling pathway in scleral fibroblasts is not known. Interestingly, recent work in mice has shown that exogenous treatment with RA can alter sclera biomechanics without noticeable changes in GAG production^11^. Our *in vitro* collagen contraction assay allows us to pick up early changes in cell behavior, particularly those linked to actin cytoskeleton dynamics and mechano-responses^24,25,68^. In turn, it is well known that such changes are the prelude to internal signaling cascades that regulate a wide range of cellular processes, including motility, metabolism, proliferation, apoptosis and differentiation^69^. A number of studies suggest that RA can affect fibroblast-mediated gel contraction, independent of its effect of transcription regulation. In pediatric gingival^70^ and Tenon’s capsule^71^ fibroblasts, RA was shown to inhibit gel contraction and triggered morphological changes similar to what we observed for scleral fibroblasts, including loss of protrusions and disruption of the actin cytoskeleton^70,71^. More recently, RARb activation was shown to inhibit hepatic stellate cells’ ability to contract collagen gel contraction through downregulation of myosin light chain 2 expression^72^. RA has also been shown to affect actin dynamics in platelets, independently of its effect on transcription, interacting directly with proteins affecting actomyosin regulation, including Rho kinase (ROCK) and myosin^73^. Similarly, in embryonic epicardial cells, RAR activation leads to cytoskeletal reorganization via the RhoA-ROCK pathway^74^, and in cancer cells and carcinoma associated fibroblasts RARβ was shown to modulate mechanosensing and invasion via myosin light chain 2^41,75^. In ocular fibroblasts, we have shown that changes in fibroblast gel contraction potential directly reflect changes in cell mechanostat that have been linked to pathologies affecting tissue structure and stiffness and extra-cellular matrix composition, including conjunctival and orbital fibrosis and Floppy Eyelid Syndrome^24–26,46^.

While more work will be needed to understand how RA affects actin dynamics in scleral fibroblasts, it is likely that this immediate response would be followed by downstream transcriptional changes leading to changes in proliferation and matrix remodeling that would modulate scleral growth and homeostasis *in vivo*^69,75^.

In conclusion, we present here a novel *in vitro* model to study human choroid and sclera stromal cell interactions. We show for the first time these two cell types can interact through chemical mediators and that these interactions are restricted by age and antero-posterior location. We present evidence for a novel mechanism by which dopamine may regulate postnatal eye growth by affecting the sclera-choroid stroma interactions through the induction of exosome-associated RA release. Our work suggests that the growing human postnatal sclera and choroid are very different from the adult mature tissues and that the retina-to-sclera signals modulating postnatal eye growth might be involving transient, post-transcriptional regulations of cellular interactions, more suited to the constant fine-tuning occurring during normal vision-led eye growth. We anticipate these signals may be different from the strong signals imposed by visual manipulations in animal models and postulate that our model will be useful to decipher cellular and molecular components of the retina-to-sclera cascade regulating human eye growth.

## METHODS

### Primary cell cultures

Donor tissue was obtained from the NHS Blood and Tissue transplant UK (https://www.nhsbt.nhs.uk/tissue-and-eye-services/<u>; donors 5, 8, 9, 14 and 15</u>) and from the NIIOS Amnitrans Eye Bank Rotterdam (http://www.niios.com/amnitrans-eyebank-rotterdam/; donor 10), with consent for research use in accordance with the tenets of the Declaration of Helsinki and Moorfields Biobank Ethics Committee approval (Moorfields Biobank Ethics Committee Reference: 20/SW/0021-2018ETR73). Donor details are shown in Table S1.

The choroid was detached from the scleral shell, and the retina removed. The sclera was cut into 4 quadrants (nasal, temporal, inferior, superior), and 3 anterior to posterior sections (anterior, equatorial, and posterior).

To generate primary choroid stroma cultures, the choroid was chopped finely with scissors, transferred to a tube with 5 ml of PBS and further mechanically dissociated by pipetting up and down with a fine pipette. The resulting suspension was centrifuged at 340g for 5 min. The pellet was resuspended in 2 ml of complete culture medium [Dulbecco’s Modified Eagle’s Medium (DMEM,4.5 g/L glucose, Gibco, Life Technologies, UK) supplemented with 10% foetal bovine serum (Sigma), penicillin (100U/ml)/streptomycin (100 ug/ml, Gibco), and L-glutamine (2mM, Gibco), and transferred to a tissue culture dish. The largest choroid fragments were covered with 20x20 mm^2^ sterile coverslips to secure them in place and ensure close contact to the bottom of the dish.

To generate primary scleral fibroblast cultures, the scleral sections were cut into small fragments (< 2x2 mm^2^) and incubated for 15-20 minutes with 0.05% Collagenase D (Roche # 1088858) in DPBS containing Ca2+ and Mg2+ (Gibco # 14040-091). The sclera fragments were then transplanted onto a 100 mm diameter tissue culture dish using forceps and covered with 20x20 mm^2^ sterile coverslips. 6 ml of complete culture medium was then added to the dish gently so as not to disturb the coverslips.

Both choroid and sclera tissues were then placed in the incubator at 37°C, 5% CO2, and allowed to grow. The medium was replaced weekly, and the tissues were checked for fibroblast growth emanating from the explants every 2 weeks. When cell growth covered more than 60% of the dish, the coverslips and larger tissue fragments were removed and the cells were trypsinized (0.05% Trypsin-EDTA, Sigma, UK) and transferred to T75 tissue culture flasks (Corning, USA). The resulting primary cultures were expanded and stock aliquots frozen at passages 2 and 3. The cells were used between passage 4 and 9 for all experiments.

### Gel Contraction Assays

The collagen contraction assay was performed as previously^24^, with minor modifications. Briefly, scleral fibroblasts were seeded in a 1.5 mg/mL collagen type-I matrix (First Link (UK) Ltd, Wolverhampton, UK) at a final concentration of 7 × 10^4^ cells/ml. 150µl of the suspension were cast into the wells of the 35 mm diameter glass bottom MatTek dishes (MatTek Corporation, USA), and the gels were allowed to polymerise in the incubator for 5-10 minutes. The gels were then detached from the edge of the well, and floated into 2 ml of medium: control serum-free DMEM (SFM), choroid conditioned medium (CCM) or dopamine-CCM, with/without addition of inhibitors as required. When inhibitors were used, the corresponding control medium was treated with an identical volume of inhibitor dilution solution. Gel contraction was monitored at days 1 and 4 by digital photography. Gel areas were measured using the open-source software Fiji (https://imagej.net/software/fiji/). Contraction was plotted as a percentage of gel area normalized to original area [contraction (%) = (1 - gel area / microwell area)x100]^24^.

To assess the effect of retinoic acid (RA; Merck/Sigma, UK) and retinoic acid receptor (RAR) antagonist (*AGN193109,* Bio-Techne Ltd./Tocris) efficacy on serum stimulated contraction, scleral fibroblasts were embedded in collagen gels as above and the gels were incubated in complete medium (DMEM 10% serum, penicillin/streptomycin, glutamine) in the presence of 1µM RA, 1µM *AGN193109* or both.

### Choroid Conditioned Medium

The standard choroid conditioned medium (CCM) was prepared by incubating sub-confluent choroid cells in T75 flasks with 8 ml of DMEM supplemented with 2mM L-glutamine (SFM) for 24 hours. The medium was then collected and filtered with a 0.45 um filter to remove cellular debris. Serum-free medium was placed in empty tissue culture T25 flasks (Corning, USA) in the incubator under the same conditions as a control. Dopamine-treated choroid conditioned medium (dopamine-CCM) was generated by treating the choroid cells with various concentrations of dopamine hydrochloride prior to preparing the conditioned medium. Briefly, a 100uM stock solution of dopamine hydrochloride (Sigma-Aldrich, UK) was prepared in water and sterile filtered. The stock solution was further diluted in complete medium to final concentrations ranging from 1nM to 10 uM. Choroid cells were cultured in complete medium with dopamine for 24 hours or 96h depending on the experiment, after which the dopamine-containing complete media was replaced with 8 ml of SFM. Following 24 hours of incubation, the resulting dopamine-CCM was collected and processed like the standard CCM. Where required, retinoic acid (1uM), retinoic acid receptor antagonist (*AGN193109, 1uM)* or both were added to the filtered CCM or dopamine-CCM immediately prior to adding the medium onto the gels for the contraction assay.

To produce exosome-depleted CCM and exosome-depleted dopamine-CCM, the collected conditioned media were centrifuged at 100,000g for 2 hours at 4°C. After ultracentrifugation, 90% of the tube contents were collected without touching the bottom of the ultracentrifuge tube and sterile filtered. For experiments comparing exosome-depleted CCM and exosome-depleted dopamine-CCM to other conditions, all other conditions were also sterile filtered for consistency.

To produce inhibitor-treated-dopamine-CCM, choroid cells were incubated with 10nM dopamine and 1µM GW4869 (Sigma-Aldrich, UK) or 8µM WIN18446 (Sigma-Aldrich, UK) in complete medium for 24 hours, after which the media was replaced with SFM to produce the CCM as above. Both inhibitors were diluted to a stock concentration 625 times greater than the desired final concentration in sterile DMSO, and a matching choroid flask was treated with the same volume of plain DMSO as vehicle control.

### Retinoic Acid ELISA

Human Retinoic Acid ELISA Kit (Biorbyt, UK, Cat# orb407080) was used to measure RA concentration following the manufacturer’s protocol. CCM was collected and filtered (0.45 filter, Sigma-Aldrich, UK) to remove cellular debris. About 5% of each sample was aliquoted out and stored at 4C until the assay was performed. The remaining CCM was centrifuged at 100,000g for 2 hours at 4°C to isolate the exosomes. After the centrifugation, all the supernatant media was removed and stored at 4°C, without disturbing the exosome pellet. The exosome pellet was lysed using 80µl RIPA buffer (50 mM Tris HCl pH8.0, 150 mM NaCl, 1% Triton-100, 0.5% Sodium deoxycholate and 0.1% SDS) and vortexed. The media samples and exosome fraction were then diluted in the kit sample diluent (1:1 for the media; 1:100 dilution for the exosome fraction). 50µl of each diluted sample were loaded on a 96 well plate in duplicate alongside the standard RA concentrations from the kit, and 50µl of HRP solution were rapidly added to each well. The plate was covered with an adhesive film and incubated at 37°C for 1 hour. The plate was then washed using the kit washing buffer 5 times for 2 minutes each. 90µl of the TMB substrate was then added and the plate was incubated at 37°C for 20 minutes. After the final incubation, 50µl of the stop solution was quickly added and the plate was analysed using the Spark® Multimode Microplate Reader (Tecan, Switzerland), measuring absorbance at 450nm, using 570nm as reference. The final RA concentration values were calculated by adjusting the absorbance values to the standard curve using the Sigmoidal 4PL function in GraphPad Prism.

### Immunofluorescence

Immunofluorescence (2D) for choroid cell markers was performed as previously^76^. Briefly, cells on coverslips were fixed with 3.7% formaldehyde in PBS for 7 minutes, and permeabilised with 0.5% Triton -X 100 (Sigma-Aldrich, UK) in PBS for 20 minutes, followed by 0.1M Glycine (Sigma-Aldrich, UK) in PBS for 10 minutes. The coverslips were transferred to a humidified dark chamber and blocked/stained for F-actin with Rhodamine-labeled Phalloidin (Molecular Probes, OR, USA) at 1:20 dilution in TBS-1% BSA-1% FBS at room temperature for 20 minutes. The coverslips were then incubated with primary antibody (Table S2) diluted in TBS-1% BSA at room temperature for 1 hour, followed by washing (3 times 10 minutes), and incubation in secondary antibody (diluted in TBS-1% BSA) at room temperature for 1 hour. Following washes, the coverslips were mounted on slides with Fluoroshield mounting medium with DAPI (Abcam, UK). Images were taken using an EVOS M7000 microscope with onboard M7000 software with 10X objective.

Immunofluorescence (3D) for scleral cells in gels was as previously published^48^. Briefly, the gels were fixed with 3.7% formaldehyde in PBS for 30 minutes, followed by 0.5% Triton -X 100 (Sigma-Aldrich, UK) permeabilization for 30 minutes, and 0.1M Glycine (Sigma-Aldrich, UK) for 30 minutes. The gels were then incubated with TRITC-conjugated phalloidin (HelloBio, UK; 1:20 in TBS with 1% BSA and 1% FBS) for F-actin staining at room temperature in the dark for 30 minutes, followed by washing with TBS/BSA pH 8.0 for 30 minutes. The gels were then incubated with anti-phospho-myosin [pMLC2] primary antibody for 1.5 hours at room temperature followed by washing (3 times; with TBS-1%BSA) for 30 minutes, and further incubation with secondary antibody diluted in TBS/BSA for 1.5 hours. The gels were washed 2 more times in TBA-1%BSA and 1 time in TBS alone before being mounted onto slides using Fluoroshield mounting medium with DAPI (Abcam, UK). Images were taken with a Leica Stellaris5 confocal microscope with 10X objective and analysed as Maximum Projection (Z-projects) in ImageJ/Fiji. For the analysis of the scleral cell morphological parameters, the gels were processed in a similar way except that FITC-conjugated Rhodamine-phalloidin (HelloBio, UK; 1:100 in TBS with 1%BSA and 1% FBS) was used to stain F-actin, and the gels were washed and mounted immediately after the phalloidin incubation. Quantification of the phospho-myosin and F-actin staining was done in ImageJ/Fidji. Threshold was applied to the Z-projections to select the cells and cells outlines identified in the F-actin images using the “Analyse particles” function, set in the ROI manager and individual integrated density was measured for each cell. The same ROIs were applied to the phospho-myosin channel. Results were expressed as mean integrated density/cell.

|  | Dilution | Supplier | Cat.No |
| --- | --- | --- | --- |
| Mouse anti- α-SMA<br>(Clone 1A4) | 1:100 | Sigma-Aldrich | A5228 |
| Rabbit anti-Calponin<br>1 antibody [EP798Y] | 1:500 | Abcam | Ab46794 |
| Rabbit anti-Vimentin<br>antibody [EPR3776] | 1:1000 | Abcam | Ab92547 |
| Rabbit anti-pMLC2<br>(Ser19) | 1:100 | Cell Signalling | 3671S |
| Alexa Fluor® 488<br>AffiniPure™ Donkey<br>Anti-Mouse IgG<br>(H+L) | 1:100 | Jackson<br>Immunoresearch<br>Laboratories, PA,<br>USA | 715-545-150 |
| Alexa Fluor® 488<br>AffiniPure™ Donkey<br>Anti-Rabbit IgG<br>(H+L) | 1:100 | Jackson<br>Immunoresearch<br>Laboratories, PA,<br>USA | 711-545-152 |

### RNA extraction, library preparation and sequencing

Subconfluent choroid cell cultures were incubated with fresh complete medium with/without 10nM dopamine for 24 hours. The medium was then replaced with 8ml serum-free DMEM supplemented with 2mM L-glutamine (SFM) and the cells placed back in the incubator for 1 hour. The cells were then trypsinised, centrifuged and RNA was extracted using the RNeasy Plus Mini Kit (Qiagen 74134 &74136) according to the manufacturer’s instruction.

Concentration/purity was evaluated using absorbance at 260/280 and 260/230 (Nanodrop ND-1000), and samples were stored at −80 °C until use. The RNA libraries were prepared using the *KAPA mRNA HyperPrep Kit for Illumina* using the NextSeq2000 platform at the UCL Genomics facilities (https://www.ucl.ac.uk/child-health/research/genetics-and-genomic-medicine/ucl-genomics).

### Bioinformatics analysis

The differential RNASeq analysis was performed using the R software, Bioconductor packages including DESeq2 and the SARTools package developed at PF2 - Institut Pasteur (doi: http://dx.doi.org/10.1371/journal.pone.0157022). Normalization and differential analysis were carried out according to the DESeq2 model and package. An fdr p-value adjustment was performed [8 and BY2001] and the level of controlled false positive rate was set to 0.05. The free online platform Galaxy (https://usegalaxy.org/) was used to create the DEGs heatmap plot, using all significant DEGs (adjusted p value <0.05). Gene Ontology (GO) function analysis and Kyoto Encyclopedia of Genes and Genomes (KEGG) pathway analysis of DEGs were performed with String (https://string-db.org/cgi/input?sessionId=bANRLvdHyc4g&input_page_show_search=off), and plotted using the platform Signal (weighted harmonic mean between the observed/expected gene ratio and -log(FDR), FRD value and Gene counts parameters).

### LDH assay

CyQuant LDH (Lactate Dehydrogenase) cytotoxicity assay kit (Invitrogen, Thermo Fisher Scientific, UK) was used to evaluate dopamine toxicity. Choroid cells were seeded onto a 96 well plate in complete medium (4,000 cells/well/100ul medium) and incubated overnight at 37°C, 5% CO_2_. The medium was then replaced with fresh medium supplemented with 0, 10nM, 100nM or 1uM Dopamine, and the cells were placed back in the incubator. After 24 and 96 hours of dopamine treatment, the culture medium was processed following the manufacturer’s protocol, and the cytotoxicity percentage was calculated using the formula: Cytotoxicity (%) = (compound-treated LDH activity – spontaneous LDH activity)/(maximum LDH activity – spontaneous LDH activity). Cells were lysed with lysis buffer as maximum LDH activity controls (100% cytotoxicity). All conditions were processed in triplicates.

### Live/Dead Viability Assay

The Live/Dead Viability assay (Life Technologies, UK) was used to determine the effect of dopamine-CCM on scleral fibroblasts in gels. Scleral fibroblasts were embedded in collagen gels as per standard contraction assay protocol, using CCM and dopamine-CCM. After 1 and 4 days, the gels were transferred from MatTek dishes into 24-well plates, and incubated with 2uM calcein AM and 4uM ethidium homodimer-1 in PBS at room temperature for 40 minutes. Images were taken using Zeiss LSM 700 confocal microscope with onboard Zen software with 10X objective.

### Cell proliferation (2D)

6,250 scleral fibroblasts were seeded into each well in a 12 well plates (Nunc, Thermo, UK) and incubated at 37°C, 5% CO2 overnight in complete medium with 1µM of RA or control medium with DMSO (diluted similarly). Duplicate wells were trypsinised and cells counted with a hemocytometer on days 1, 3 and 7.

### Cell proliferation (3D)

Scleral fibroblasts were embedded in collagen gels as per standard contraction assay and incubated with CCM or dopamine-CCM. After 1 day and 4 days, the gels were transferred from the MatTek dishes to a 48-well plate, with 100ul of matched conditioned medium. 10ul CCK8 reagent (Cell Counting Kit-8, Sigma-Aldrich, UK) was added to the wells and the gels were further incubated for 2 hours. 100ul of the medium was then transferred to a 96-well plate, and absorbance was measured at 450nm (Tecan Safire). Samples were processed as duplicates. In a parallel set, the gels were digested with 0.05% Collagenase D (Roche # 1088858) in DPBS (containing Ca2+ and Mg2+) (Gibco # 14040-091) on a shaking incubator at 37°C for 5-15 mins until dissolved^40^. The enzyme was diluted by addition of 3ml of PBS and the suspension was centrifuged 340 g for 7 min to pellet the cells. The pellet was resuspended in PBS and cells counted with a hemocytometer.

The effect of retinoic acid on baseline scleral cell metabolic activity/proliferation in 3D gels was measured using Alamar Blue as previously^47^. Scleral fibroblasts were embedded in collagen gels and allowed to contract in complete medium at 37°C, 5% CO2. At days 1, 3 and 7, 200µl (10% of the medium volume) of Alamar Blue reagent (Thermo Fisher Scientific) was added and the dishes were placed back in the incubator for 2 hours. 100µl of the medium was then taken from each sample and transferred in triplicates to a 96-well plate.

The fluorescence intensity was measured at 530 of excitation per 590 of emission using a plate reader (Tecan Safire).

### Statistical analysis

Data in figures and text are presented as mean ± standard error of the mean (SEM) for a minimum of 3 independent repeats unless otherwise stated. Statistical tests were performed using GraphPad Prism 10 software. Statistical significance was determined using paired Student’s t-tests and one- or two-way analysis of variance (ANOVA). P-values ≤ 0.05 were considered statistically significant for all experiments.

## ACKNOWLEDGMENTS

This study was supported by grants from Moorfields Eye Charity (R180011A to M.B.), Sight Research UK (SAC053 to M.B.), UCLTherapeutic Acceleration Support (TAS) Fund/ MOORFIELDS EYE HOSPITAL NHS FOUNDATION TRUST (BRC4-02-RB401-105 to M.B.). ADN is supported by the National Institute for Health Research (NIHR) Moorfields Biomedical Research Centre. This report is independent research. NHS Blood & Transplant have provided material in support of the research. The views expressed in this publication are those of the author(s) and not necessarily those of NHS Blood & Transplant. Tissue for this project was provided through Moorfields Biobank and supported by NIHR funding to Prof. Marcus Fruttiger. We are particularly grateful to the families who have consented to donate paediatric tissue.

---

**Table S1.** Donor details.

| Donor ID | Sex | Age | Eye | Cause of death | DTP | Primary |
| --- | --- | --- | --- | --- | --- | --- |

|  |  |  |  |  | (days) | cells used<br>in this<br>study |
| --- | --- | --- | --- | --- | --- | --- |
| Donor 5 | Male | 42 | OD | unavailable | 4 | Choro5R |
| Donor 8 | Male | 31 | OD | Subarachnoid<br>Haemorrhage | 5 | Choro8R |
| Donor 9 | Male | 3 | OD | Cardiac Arrest | 3 | Choro9R<br>HSF9 |
| Donor 10 | Female | 35 | OD | Septic shock | 4 | Choro10R |
| Donor 14 | Female | 3 | OS | Pleuropulmonary<br>blastoma Type 2 | 3 | Choro14L<br>HSF14 |
| Donor 15 | Male | 7 | OS | Sudden<br>unexpected death | 3 | Choro15L |

\*DTP, death to processing time

## REFERENCES

1. Klaver C, Polling JR, Group EMR. Myopia management in the Netherlands. Ophthalmic Physiol Opt. Mar 2020;40(2):230–240. doi:10.1111/opo.12676

2. Harper AR, Summers JA. The dynamic sclera: extracellular matrix remodeling in normal ocular growth and myopia development. Exp Eye Res. Apr 2015;133:100–11. doi:10.1016/j.exer.2014.07.015

3. Metlapally R, Wildsoet CF. Scleral Mechanisms Underlying Ocular Growth and Myopia. Prog Mol Biol Transl Sci. 2015;134:241–8. doi:10.1016/bs.pmbts.2015.05.005

4. Zhang Y, Wildsoet CF. RPE and Choroid Mechanisms Underlying Ocular Growth and Myopia. Prog Mol Biol Transl Sci. 2015;134:221–40. doi:10.1016/bs.pmbts.2015.06.014

5. Summers JA, Cano EM, Kaser-Eichberger A, Schroedl F. Retinoic acid synthesis by a population of choroidal stromal cells. Exp Eye Res. 12 2020;201:108252. doi:10.1016/j.exer.2020.108252

6. Summers JA, Schaeffel F, Marcos S, Wu H, Tkatchenko AV. Functional integration of eye tissues and refractive eye development: Mechanisms and pathways. Exp Eye Res. 08 2021;209:108693. doi:10.1016/j.exer.2021.108693

7. Ostrin LA, Harb E, Nickla DL, et al. IMI-The Dynamic Choroid: New Insights, Challenges, and Potential Significance for Human Myopia. Invest Ophthalmol Vis Sci. May 01 2023;64(6):4. doi:10.1167/iovs.64.6.4

8. Shelton L, Troilo D, Lerner MR, Gusev Y, Brackett DJ, Rada JS. Microarray analysis of choroid/RPE gene expression in marmoset eyes undergoing changes in ocular growth and refraction. Mol Vis. Aug 2008;14:1465–79.

9. Brown DM, Mazade R, Clarkson-Townsend D, Hogan K, Datta Roy PM, Pardue MT. Candidate pathways for retina to scleral signaling in refractive eye growth. Exp Eye Res. Jun 2022;219:109071. doi:10.1016/j.exer.2022.109071

10. Huang L, Ye L, Li R, et al. Dynamic human retinal pigment epithelium (RPE) and choroid architecture based on single-cell transcriptomic landscape analysis. Genes Dis. Nov 2023;10(6):2540–2556. doi:10.1016/j.gendis.2022.11.007

11. Brown DM, Yu J, Kumar P, et al. Exogenous All-Trans Retinoic Acid Induces Myopia and Alters Scleral Biomechanics in Mice. Invest Ophthalmol Vis Sci. May 1 2023;64(5):22. doi:10.1167/iovs.64.5.22

12. McFadden SA, Howlett MH, Mertz JR, Wallman J. Acute effects of dietary retinoic acid on ocular components in the growing chick. Exp Eye Res. Oct 2006;83(4):949–61. doi:10.1016/j.exer.2006.05.002

13. Mertz JR, Wallman J. Choroidal retinoic acid synthesis: a possible mediator between refractive error and compensatory eye growth. Exp Eye Res. Apr 2000;70(4):519–27. doi:10.1006/exer.1999.0813

14. Rada JA, Palmer L. Choroidal regulation of scleral glycosaminoglycan synthesis during recovery from induced myopia. Invest Ophthalmol Vis Sci. Jul 2007;48(7):2957–66. doi:10.1167/iovs.06-1051

15. Rada JA, Hollaway LR, Lam W, Li N, Napoli JL. Identification of RALDH2 as a visually regulated retinoic acid synthesizing enzyme in the chick choroid. Invest Ophthalmol Vis Sci. Mar 2012;53(3):1649–62. doi:10.1167/iovs.11-8444

16. Marzani D, Wallman J. Growth of the two layers of the chick sclera is modulated reciprocally by visual conditions. Invest Ophthalmol Vis Sci. Aug 1997;38(9):1726–39.

17. McFadden SA, Howlett MH, Mertz JR. Retinoic acid signals the direction of ocular elongation in the guinea pig eye. Vision Res. Mar 2004;44(7):643–53. doi:10.1016/j.visres.2003.11.002

18. Read SA, Alonso-Caneiro D, Vincent SJ, et al. Anterior eye tissue morphology: Scleral and conjunctival thickness in children and young adults. Sci Rep. 09 2016;6:33796. doi:10.1038/srep33796

19. Summers JA, Jones KL. Single Cell Transcriptomics Identifies Distinct Choroid Cell Populations Involved in Visually Guided Eye Growth. Front Ophthalmol (Lausanne). 2023;3doi:10.3389/fopht.2023.1245891

20. Lehmann GL, Hanke-Gogokhia C, Hu Y, et al. Single-cell profiling reveals an endothelium-mediated immunomodulatory pathway in the eye choroid. J Exp Med. Jun 01 2020;217(6)doi:10.1084/jem.20190730

21. Voigt AP, Whitmore SS, Mulfaul K, et al. Bulk and single-cell gene expression analyses reveal aging human choriocapillaris has pro-inflammatory phenotype. Microvasc Res. 09 2020;131:104031. doi:10.1016/j.mvr.2020.104031

22. Haddad A, Laicine EM, Tripathi BJ, Tripathi RC. An extensive system of extravascular smooth muscle cells exists in the choroid of the rabbit eye. Exp Eye Res. Sep 2001;73(3):345–53. doi:10.1006/exer.2001.1042

23. May CA. Non-vascular smooth muscle cells in the human choroid: distribution, development and further characterization. J Anat. Oct 2005;207(4):381–90. doi:10.1111/j.1469-7580.2005.00460.x

24. Dahlmann-Noor AH, Martin-Martin B, Eastwood M, Khaw PT, Bailly M. Dynamic protrusive cell behaviour generates force and drives early matrix contraction by fibroblasts. Exp Cell Res. Dec 10 2007;313(20):4158–69.

25. Ezra D, Ellis J, Beaconsfield M, Collin R, Bailly M. Changes in Fibroblast Mechanostat Set Point and Mechanosensitivity: An Adaptive Response to Mechanical Stress in Floppy Eyelid Syndrome. Investigative Ophthalmology & Visual Science. 2010 2010;51(8)(0146-0404):3853-3863. doi:10.1167/iovs.09-4724

26. Kechagia JZ, Ezra DG, Burton MJ, Bailly M. Fibroblasts profiling in scarring trachoma identifies IL-6 as a functional component of a fibroblast-macrophage pro-fibrotic and pro-inflammatory feedback loop. Scientific reports. 2016;6:28261. doi:10.1038/srep28261 10.1038/srep28261.

27. Wallman J, Winawer J. Homeostasis of eye growth and the question of myopia. Neuron. Aug 19 2004;43(4):447–68. doi:10.1016/j.neuron.2004.08.008

28. Avetisov ES, Savitskaya NF, Vinetskaya MI, Iomdina EN. A study of biochemical and biomechanical qualities of normal and myopic eye sclera in humans of different age groups. Metab Pediatr Syst Ophthalmol. 1983;7(4):183–8.

29. Ming M, Li X, Fan X, et al. Retinal pigment epithelial cells secrete neurotrophic factors and synthesize dopamine: possible contribution to therapeutic effects of RPE cell transplantation in Parkinson’s disease. J Transl Med. Jun 28 2009;7:53. doi:10.1186/1479-5876-7-53

30. Zhou X, Pardue MT, Iuvone PM, Qu J. Dopamine signaling and myopia development: What are the key challenges. Prog Retin Eye Res. Nov 2017;61:60–71. doi:10.1016/j.preteyeres.2017.06.003

31. Mathis U, Feldkaemper M, Liu H, Schaeffel F. Studies on the interactions of retinal dopamine with choroidal thickness in the chicken. Graefes Arch Clin Exp Ophthalmol. Feb 2023;261(2):409–425. doi:10.1007/s00417-022-05837-w

32. Monavarfeshani A, Yan W, Pappas C, et al. Transcriptomic analysis of the ocular posterior segment completes a cell atlas of the human eye. Proc Natl Acad Sci U S A. Aug 22 2023;120(34):e2306153120. doi:10.1073/pnas.2306153120

33. Collin J, Hasoon MSR, Zerti D, et al. Single-cell RNA sequencing reveals transcriptional changes of human choroidal and retinal pigment epithelium cells during fetal development, in healthy adult and intermediate age-related macular degeneration. Hum Mol Genet. May 5 2023;32(10):1698–1710. doi:10.1093/hmg/ddad007

34. Alexander N, Walshe J, Richardson NA, et al. Stromal cells cultivated from the choroid of human eyes display a mesenchymal stromal cell (MSC) phenotype and inhibit the proliferation of choroidal vascular endothelial cells in vitro. Exp Eye Res. Nov 2020;200:108201. doi:10.1016/j.exer.2020.108201

35. David O, Barrera I, Gould N, Gal-Ben-Ari S, Rosenblum K. D1 Dopamine Receptor Activation Induces Neuronal eEF2 Pathway-Dependent Protein Synthesis. Front Mol Neurosci. 2020;13:67. doi:10.3389/fnmol.2020.00067

36. Zimbelman AR, Wong B, Murray CH, Wolf ME, Stefanik MT. Dopamine D1 and NMDA Receptor Co-Regulation of Protein Translation in Cultured Nucleus Accumbens Neurons. Neurochem Res. Nov 21 2024;50(1):27. doi:10.1007/s11064-024-04283-w

37. Fischer AJ, Wallman J, Mertz JR, Stell WK. Localization of retinoid binding proteins, retinoid receptors, and retinaldehyde dehydrogenase in the chick eye. J Neurocytol. Jul 1999;28(7):597–609. doi:10.1023/a:1007071406746

38. Li C, McFadden SA, Morgan I, et al. All-trans retinoic acid regulates the expression of the extracellular matrix protein fibulin-1 in the guinea pig sclera and human scleral fibroblasts. Mol Vis. Apr 15 2010;16:689–97.

39. Goncalves MB, Wu Y, Clarke E, et al. Regulation of Myelination by Exosome Associated Retinoic Acid Release from NG2-Positive Cells. J Neurosci. Apr 17 2019;39(16):3013–3027. doi:10.1523/JNEUROSCI.2922-18.2019

40. Martin-Martin B, Tovell V, Dahlmann-Noor AH, Khaw PT, Bailly M. The effect of MMP inhibitor GM6001 on early fibroblast-mediated collagen matrix contraction is correlated to a decrease in cell protrusive activity. Eur J Cell Biol. Jan 2011;90(1):26–36. doi:S0171-9335(10)00199-8 [pii] 10.1016/j.ejcb.2010.09.008

41. Matellan C, Lachowski D, Cortes E, et al. Retinoic acid receptor β modulates mechanosensing and invasion in pancreatic cancer cells via myosin light chain 2. Oncogenesis. May 02 2023;12(1):23. doi:10.1038/s41389-023-00467-1

42. Hu A, Yang Y, Zhang S, Zhou Q, Wei W, Wang Y. 4-Amino-2-trifluoromethyl-phenyl retinate inhibits the migration of BGC-823 human gastric cancer cells by downregulating the phosphorylation level of MLC II. Oncol Rep. Oct 2014;32(4):1473–80. doi:10.3892/or.2014.3343

43. Troilo D, Smith EL, Nickla DL, et al. IMI - Report on Experimental Models of Emmetropization and Myopia. Invest Ophthalmol Vis Sci. Feb 2019;60(3):M31–M88. doi:10.1167/iovs.18-25967

44. Palumaa T, Balamurugan S, Pardue MT. Meta-analysis of retinal transcriptome profiling studies in animal models of myopia. Front Med (Lausanne). 2024;11:1479891. doi:10.3389/fmed.2024.1479891

45. LeBleu VS, Neilson EG. Origin and functional heterogeneity of fibroblasts. FASEB J. 03 2020;34(3):3519–3536. doi:10.1096/fj.201903188R

46. Li H, Fitchett C, Kozdon K, et al. Independent adipogenic and contractile properties of fibroblasts in graves’ orbitopathy: an in vitro model for the evaluation of treatments. PLoS One. 2014;9(4):e95586. doi:10.1371/journal.pone.0095586 10.1371/journal.pone.0095586. eCollection 2014.

47. Kozdon K, Caridi B, Duru I, Ezra DG, Phillips JB, Bailly M. A Tenon’s capsule/bulbar conjunctiva interface biomimetic to model fibrosis and local drug delivery. PLoS One. 2020;15(11):e0241569. doi:10.1371/journal.pone.0241569

48. Yang IH, Rose GE, Ezra DG, Bailly M. Macrophages promote a profibrotic phenotype in orbital fibroblasts through increased hyaluronic acid production and cell contractility. Sci Rep. 07 2019;9(1):9622. doi:10.1038/s41598-019-46075-1

49. Seko Y, Azuma N, Yokoi T, et al. Anteroposterior Patterning of Gene Expression in the Human Infant Sclera: Chondrogenic Potential and Wnt Signaling. Curr Eye Res. 01 2017;42(1):145–154. doi:10.3109/02713683.2016.1143015

50. Xi LY, Yip SP, Shan SW, Summers-Rada J, Kee CS. Region-specific differential corneal and scleral mRNA expressions of MMP2, TIMP2, and TGFB2 in highly myopic-astigmatic chicks. Sci Rep. 09 2017;7(1):11423. doi:10.1038/s41598-017-08765-6

51. Geraghty B, Jones SW, Rama P, Akhtar R, Elsheikh A. Age-related variations in the biomechanical properties of human sclera. J Mech Behav Biomed Mater. Dec 2012;16:181–91. doi:10.1016/j.jmbbm.2012.10.011

52. Boote C, Sigal IA, Grytz R, Hua Y, Nguyen TD, Girard MJA. Scleral structure and biomechanics. Prog Retin Eye Res. 01 2020;74:100773. doi:10.1016/j.preteyeres.2019.100773

53. Panda A, Yadav A, Yeerna H, et al. Tissue- and development-stage-specific mRNA and heterogeneous CNV signatures of human ribosomal proteins in normal and cancer samples. Nucleic Acids Res. Jul 27 2020;48(13):7079–7098. doi:10.1093/nar/gkaa485

54. Feldkaemper M, Schaeffel F. An updated view on the role of dopamine in myopia. Exp Eye Res. Sep 2013;114:106–19. doi:10.1016/j.exer.2013.02.007

55. Talwar S, Mazade R, Bentley-Ford M, et al. Modulation of All-Trans Retinoic Acid by Light and Dopamine in the Murine Eye. Invest Ophthalmol Vis Sci. Mar 03 2025;66(3):37. doi:10.1167/iovs.66.3.37

56. Allen RS, Khayat CT, Feola AJ, et al. Diabetic rats with high levels of endogenous dopamine do not show retinal vascular pathology. Front Neurosci. 2023;17:1125784. doi:10.3389/fnins.2023.1125784

57. Kos MZ, Duan J, Sanders AR, et al. Dopamine perturbation of gene co-expression networks reveals differential response in schizophrenia for translational machinery. Transl Psychiatry. Dec 13 2018;8(1):278. doi:10.1038/s41398-018-0325-1

58. Smith WB, Starck SR, Roberts RW, Schuman EM. Dopaminergic stimulation of local protein synthesis enhances surface expression of GluR1 and synaptic transmission in hippocampal neurons. Neuron. Mar 03 2005;45(5):765–79. doi:10.1016/j.neuron.2005.01.015

59. Simon P, Feldkaemper M, Bitzer M, Ohngemach S, Schaeffel F. Early transcriptional changes of retinal and choroidal TGFbeta-2, RALDH-2, and ZENK following imposed positive and negative defocus in chickens. Mol Vis. Aug 24 2004;10:588–97.

60. Bitzer M, Feldkaemper M, Schaeffel F. Visually induced changes in components of the retinoic acid system in fundal layers of the chick. Exp Eye Res. Jan 2000;70(1):97–106. doi:10.1006/exer.1999.0762

61. Harper AR, Wiechmann AF, Moiseyev G, Ma JX, Summers JA. Identification of active retinaldehyde dehydrogenase isoforms in the postnatal human eye. PLoS One. 2015;10(3):e0122008. doi:10.1371/journal.pone.0122008

62. Dvoriashyna M, Bentley-Ford M, Yu J, et al. All-. bioRxiv. Feb 08 2025;doi:10.1101/2025.02.05.636685

63. Goncalves MB, Wu Y, Trigo D, et al. Retinoic acid synthesis by NG2 expressing cells promotes a permissive environment for axonal outgrowth. Neurobiol Dis. Mar 2018;111:70–79. doi:10.1016/j.nbd.2017.12.016

64. Chatterjee A, Singh R. Extracellular vesicles: an emerging player in retinal homeostasis. Front Cell Dev Biol. 2023;11:1059141. doi:10.3389/fcell.2023.1059141

65. Klingeborn M, Stamer WD, Bowes Rickman C. Polarized Exosome Release from the Retinal Pigmented Epithelium. Adv Exp Med Biol. 2018;1074:539–544. doi:10.1007/978-3-319-75402-4_65

66. Edwards MM, McLeod DS, Grebe R, et al. Glial remodeling and choroidal vascular pathology in eyes from two donors with Choroideremia. Front Ophthalmol (Lausanne). 2022;2:994566. doi:10.3389/fopht.2022.994566

67. Platzl C, Kaser-Eichberger A, Trost A, et al. Melanopsin in the human and chicken choroid. Exp Eye Res. Oct 2024;247:110053. doi:10.1016/j.exer.2024.110053

68. McGee KM, Vartiainen MK, Khaw PT, Treisman R, Bailly M. Nuclear transport of the serum response factor coactivator MRTF-A is downregulated at tensional homeostasis. Research Support, Non-U.S. Gov’t. EMBO Reports. Sep 2011;12(9):963–70. doi:10.1038/embor.2011.141 10.1038/embor.2011.141.

69. Hayward MK, Muncie JM, Weaver VM. Tissue mechanics in stem cell fate, development, and cancer. Dev Cell. Jul 12 2021;56(13):1833–1847. doi:10.1016/j.devcel.2021.05.011

70. Lew HS, Tsuchihashi M, Tenshin S, Kawata T. Effect of retinoic acid on contraction of collagen gel induced by fibroblasts. Biochem Biophys Res Commun. Nov 30 1994;205(1):455–9. doi:10.1006/bbrc.1994.2687

71. Liu Y, Kimura K, Orita T, Teranishi S, Suzuki K, Sonoda KH. Inhibition by all-trans-retinoic acid of transforming growth factor-β-induced collagen gel contraction mediated by human tenon fibroblasts. Invest Ophthalmol Vis Sci. Jun 03 2014;55(7):4199–205. doi:10.1167/iovs.13-13572

72. Cortes E, Lachowski D, Rice A, et al. Retinoic Acid Receptor-β Is Downregulated in Hepatocellular Carcinoma and Cirrhosis and Its Expression Inhibits Myosin-Driven Activation and Durotaxis in Hepatic Stellate Cells. Hepatology. Feb 2019;69(2):785–802. doi:10.1002/hep.30193

73. Rondina MT, Freitag M, Pluthero FG, et al. Non-genomic activities of retinoic acid receptor alpha control actin cytoskeletal events in human platelets. J Thromb Haemost. May 2016;14(5):1082–94. doi:10.1111/jth.13281

74. Wang S, Yu J, Jones JW, et al. Retinoic acid signaling promotes the cytoskeletal rearrangement of embryonic epicardial cells. FASEB J. Jul 2018;32(7):3765–3781. doi:10.1096/fj.201701038R

75. Chronopoulos A, Robinson B, Sarper M, et al. ATRA mechanically reprograms pancreatic stellate cells to suppress matrix remodelling and inhibit cancer cell invasion. Nat Commun. Sep 07 2016;7:12630. doi:10.1038/ncomms12630

76. Kozdon K, Fitchett C, Rose GE, Ezra DG, Bailly M. Mesenchymal Stem Cell-Like Properties of Orbital Fibroblasts in Graves’ Orbitopathy. Research Support, Non-U.S. Gov’t. Investigative ophthalmology & visual science. Sep 2015;56(10):5743–50. doi:10.1167/iovs.15-16580 10.1167/iovs.15-16580.

