## Supporting information for "Dopamine selectively regulates pediatric sclera/choroid interactions through the stimulation of exosome-associated retinoic acid"

### TITLE

### Choro9R (Paediatric Choroid)

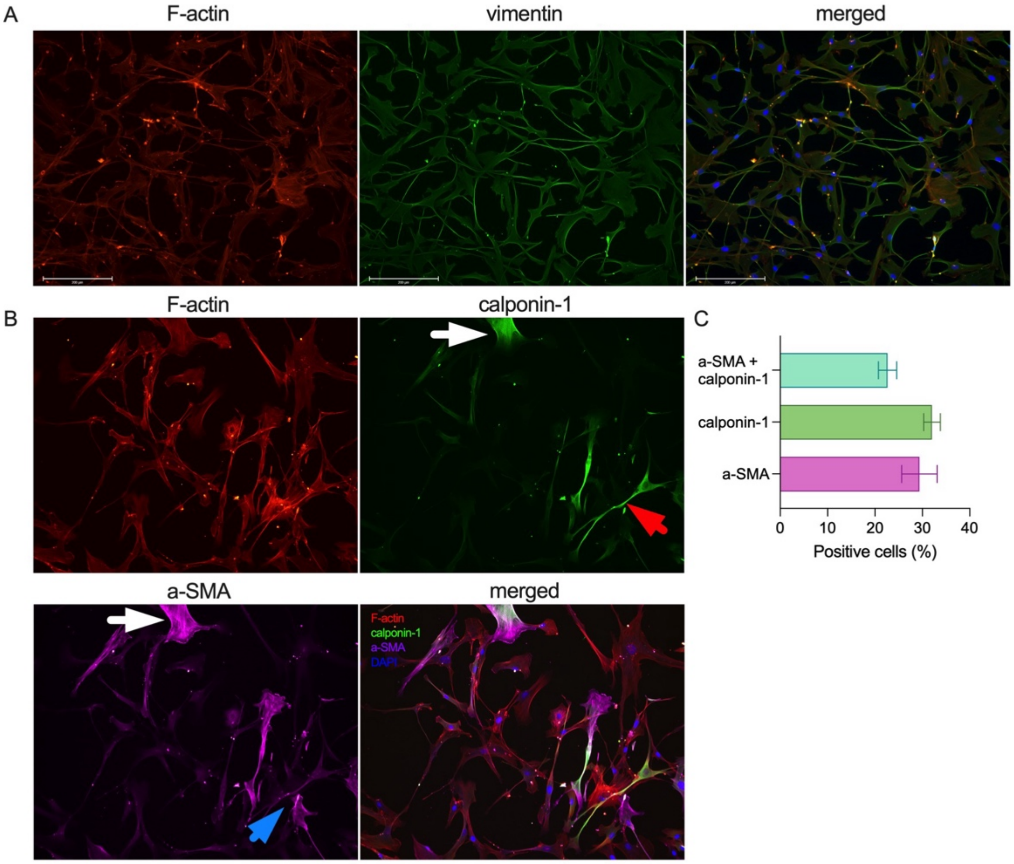


### Choro10R (Adult Choroid)

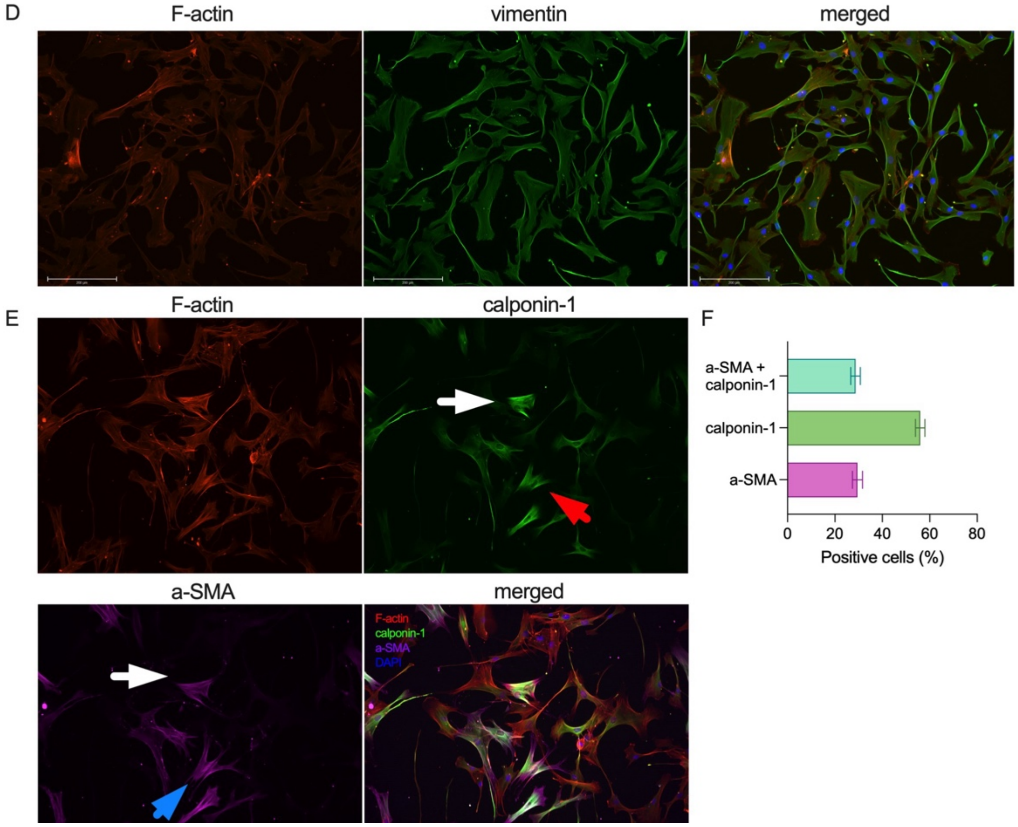

**Figure S1: Representative immunofluorescence for paediatric Choro9R *(A-C)* and adult *(D-F)* choroid cells - fibroblast and non-vascular smooth muscle cell markers.** *(A,D)* Vimentin. *(B, E)* a-SMA (blue arrow), calponin-1 (red arrow), and colocalisation of both markers (white arrow). Scale bar, 200 um. *(C,F)* Quantification of a-SMA, calponin-1, and double positive cells. Shown is percent of positive cells (mean +/- SEM, n=20).

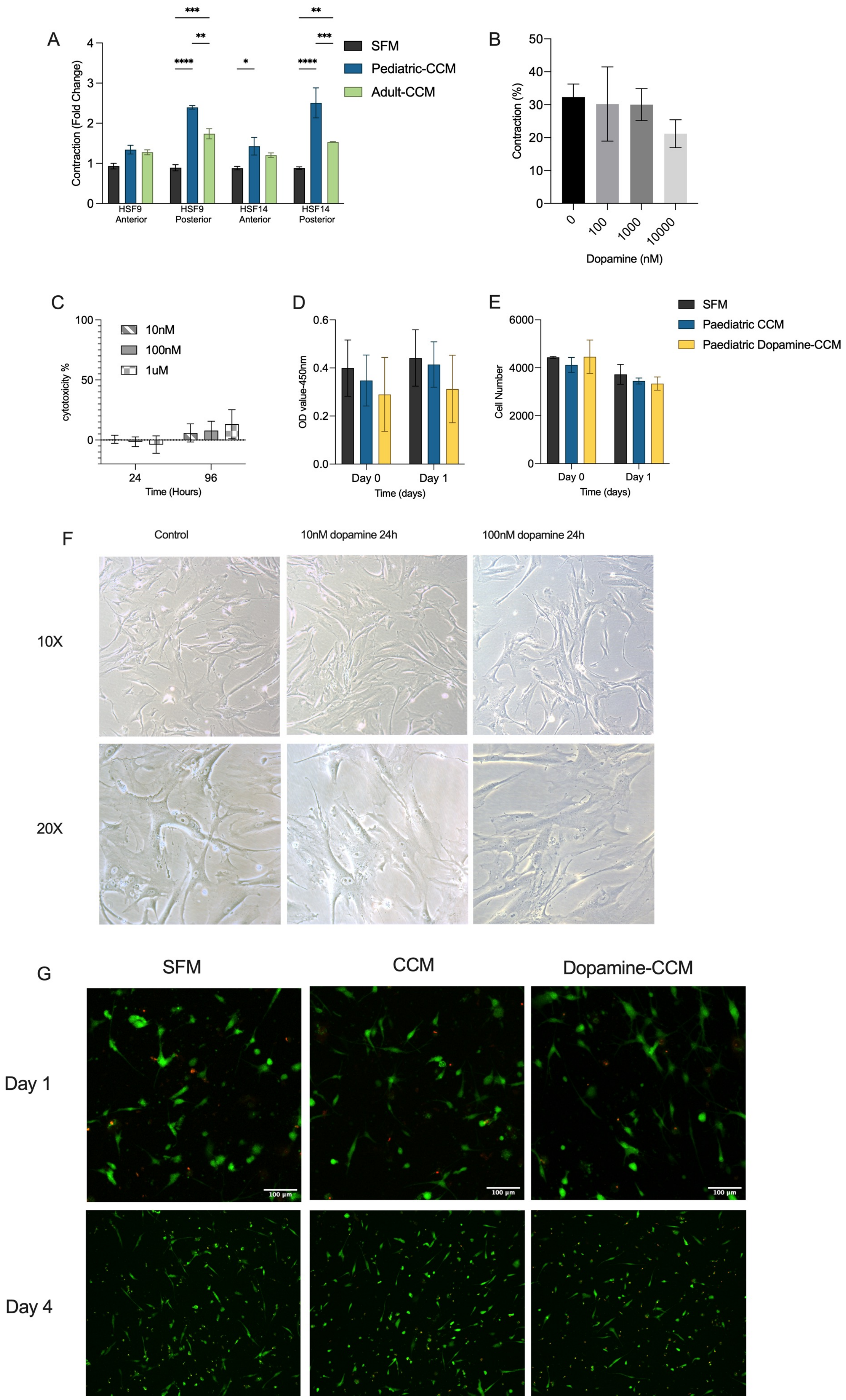
A
Contraction (Fold Change)
HSF9 Anterior
HSF9 Posterior
HSF14 Anterior
HSF14 Posterior
SFM
Pediatric-CCM
Adult-CCM
B
Contraction (%)
Dopamine (nM)
C
cytotoxicity %
Time (Hours)
10nM
100nM
1uM
D
OD value-450nm
Time (days)
E
Cell Number
Time (days)
SFM
Paediatric CCM
Paediatric Dopamine-CCM
F
Control
10nM dopamine 24h
100nM dopamine 24h
10X
20X
G
SFM
CCM
Dopamine-CCM
Day 1
Day 4

**Figure S2. Effect of dopamine and dopamine-CCM on cell viability.** *(A)* Scleral fibroblasts from anterior and posterior sclera from two paediatric donors were embedded in collagen gels and contraction was measured after 1 day in the presence of the serum-free medium (SFM) or choroid-conditioned medium (CCM) prepared from paediatric (Choro-9R; Pediatric-CCM) or adult (Choro-10R; Adult-CCM) choroid cells. Shown is mean fold change in contraction (compared to SFM) +/- SEM (SFM, n=6; each CCM, n=3). * $P<0.05$, ** $P<0.01$, *** $P<0.001$, as per Two-way ANOVA and Tukey's multiple comparisons tests. *(B)* Adult choroid cells (Choro-10R) were treated with dopamine at different concentrations (0-10 uM) for 96h, and the medium was then replaced with the serum-free medium for 24 hours to generate CCMs. These were used to stimulate paediatric scleral fibroblasts (HSF9-Posterior) contraction. Shown is mean contraction (%) +/- SEM (n>=3). No significant differences (One Way ANOVA). *(C)* Paediatric choroid cells (Choro-9R) were treated with dopamine at different concentrations for 24 or 96 hours, and cell death was measured using an LDH assay. Cytotoxicity was normalized to the 100% death control (lysed cells). Shown is mean +/- SEM, n=3. *(D, E, G)* Paediatric scleral fibroblasts (HSF9-posterior) were embedded in collagen gels and exposed to SFM, CCM or dopamine-CCM prepared from isogenic choroid cells (Choro-9L). Metabolic activity/proliferation assay (*D*, CCK8 assay) and cell numbers counts (*E*, direct cell counts) were performed at day 0 (immediately after gel preparation) and at day 1. Shown is mean +/-SEM, n>=4. No significant difference. (*G)* Representative images of Live/Dead assay on scleral fibroblasts in gels following 1 and 4 days of incubation with the different CCM. Only a few dead cells (red) could be observed (<10%) and there was no difference between the 3 different CCMs.

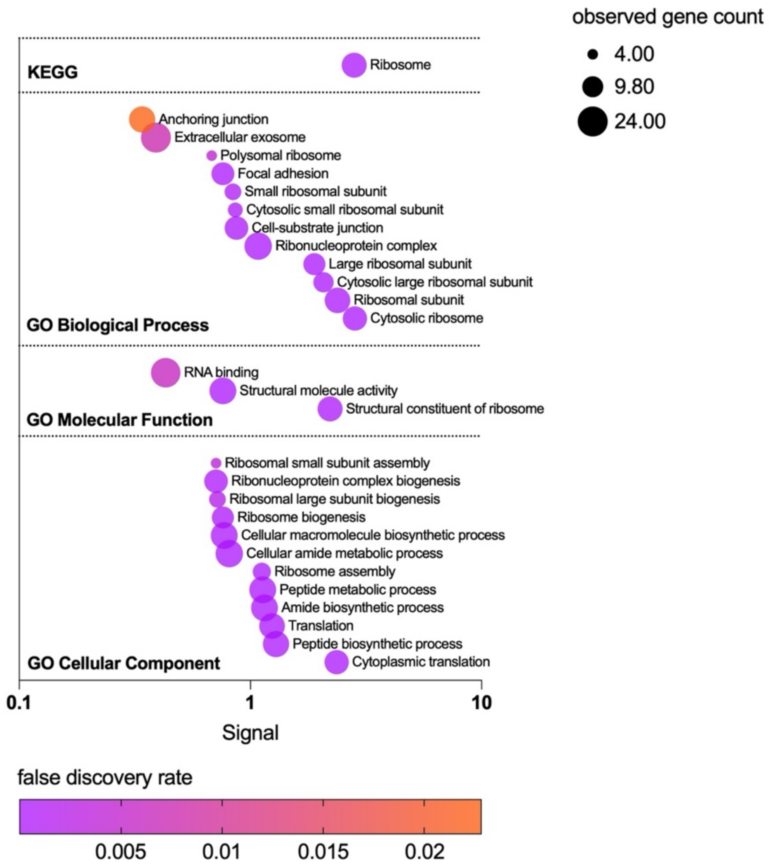


**Figure S3. Pathway enrichment** in **dopamine-treated** paediatric choroid cells compared to dopamine-treated adult choroid cells as per KEGG and Gene Ontology pathway enrichment analyses. Where the signal (observed genes ratioed to background genes and normalised to false discovery rate) is represented on the x-axis, raw observed count is represented by the point size, and the false discovery rate is represented by the point colour.

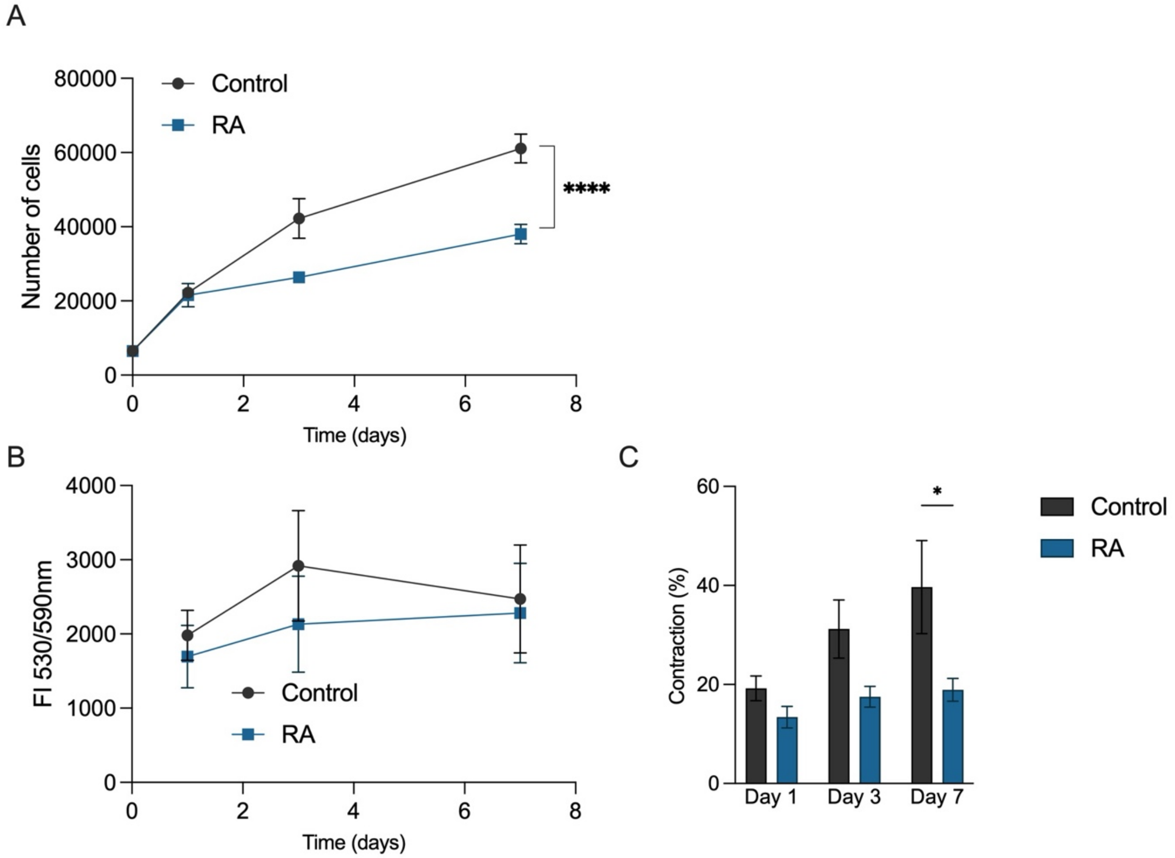


**Figure S4. Retinoic acid effects on paediatric scleral cell proliferation.** *(A)* Paediatric scleral fibroblasts (HSF9-posterior) proliferation on standard 2D substrate, in regular culture medium (10% serum) with/without 1 uM RA. Shown is the average cell number/well +/- SEM, n=3. *(B)* HS9-posterior cell proliferation/metabolic activity in collagen gels in standard culture medium (10% serum) with/without 1 uM RA, as measured with Alamar Blue. Shown is average absorbance reading/gel +/- SEM, n=3. *(C)* Matching gel contraction. Shown is mean gel contraction +/-SEM, n=3. In all experiments, the control cells were treated with an equivalent volume of solvent (DMSO) to match the RA treatment.
